# Cumulative growth and stress responses to the 2018–2019 drought in a European floodplain forest

**DOI:** 10.1101/2021.03.05.434090

**Authors:** Florian Schnabel, Sarah Purrucker, Lara Schmitt, Rolf A. Engelmann, Anja Kahl, Ronny Richter, Carolin Seele-Dilbat, Georgios Skiadaresis, Christian Wirth

## Abstract

Droughts increasingly threaten the world’s forests and their potential to mitigate climate change. In 2018–2019, Central European forests were hit by two consecutive hotter drought years, an unprecedented phenomenon that is likely to occur more frequently with climate change. Here, we examine tree growth and physiological stress responses (increase in carbon isotope composition; Δδ^13^C) to this consecutive drought based on tree-rings of dominant tree species in a Central European floodplain forest. Tree growth was not reduced for most species in 2018, indicating that water supply in floodplain forests can partly buffer meteorological water deficits. Drought stress responses in 2018 were comparable to former single drought years but the hotter drought in 2018 induced drought legacies in tree growth while former droughts did not. We observed strong decreases in tree growth and increases in Δδ^13^C across all tree species in 2019, which are likely driven by the cumulative stress both consecutive hotter droughts exerted. Our results show that consecutive hotter droughts pose a novel threat to forests under climate change, even in forest ecosystems with comparably high levels of water supply.

## Introduction

The frequency and intensity of droughts and corresponding surges of forest dieback events around the globe are projected to increase in the 21^st^ century (Allen et al., 2010; IPCC, 2014). This critically endangers the world’s forests and the variety of ecosystem services they sustain, such as their potential to act as carbon sink (Anderegg et al., 2020) and as a nature-based solution for climate change mitigation (Griscom et al., 2017). Recent drought events, moreover, belong to a new category, so called ‘hotter droughts’, where low precipitation coincides with heat waves, which creates a positive feedback loop between soil-water depletion through evapotranspiration and increased surface temperatures through reduced cooling by latent heat production (Allen, Breshears, & McDowell, 2015; Buras, Rammig, & Zang, 2020). In 2018–2019, Central Europe has been hit by two consecutive and hotter summer drought events, a phenomenon unprecedented at least in the last 250 years but likely to occur more frequently with intensifying climate change (Hari, Rakovec, Markonis, Hanel, & Kumar, 2020). The 2018 hotter drought alone had already stronger negative effects on European ecosystems than the formerly severest drought event in 2003 (Buras et al., 2020) and induced widespread premature leaf senescence and tree mortality (Schuldt et al., 2020). An increasing number of studies has shown that droughts can affect tree growth and hence carbon cycling in forests for years after the actual drought event and that such ‘legacy effects’ are widespread in forest ecosystems (e.g. Anderegg et al., 2015; Kannenberg et al., 2019; Szejner, Belmecheri, Ehleringer, & Monson, 2020). The consecutive hotter drought in 2019 may thus have critically amplified drought stress as trees were hit that already had emptied carbon reserves, impaired hydraulic functioning due to embolism and weakened defense systems (Anderegg et al., 2013; Schuldt et al., 2020) and only access to emptied soil water reserves.

Drought effects on forests can be analyzed retrospectively through analyses of tree-rings, which are an archive of past growing conditions including climate and water availability (Schweingruber, 1996). In dendroecology the annual growth of trees (i.e. the width of tree rings formed each year) is a principal indicator of drought effects, which can be analyzed through comparing growth in drought years with mean growth in a reference period, i.e. years with ‘normal’ climate conditions prior to the drought event (with growth reductions indicating drought stress) (Lloret, Keeling, & Sala, 2011; Schwarz et al., 2020). This growth response to drought can be quantified using the growth resistance index of Lloret et al. (2011), which may be an especially suitable approach when rapid impact assessments are needed and no data is available on the post disturbance period. Next to growth, the carbon isotope ratio of ^13^C to ^12^C in wood — called δ^13^C — is a widely-used physiological indicator of a treès water status and drought stress (Farquhar, Ehleringer, & Hubick, 1989; Grossiord et al., 2014; Jucker et al., 2017). Under ample water supply and fully open stomata, trees discriminate against the heavier ^13^C in favor of the lighter ^12^C. However, under water shortage, stomatal conductance is more strongly down-regulated than CO2 assimilation, which induces an increase of δ^13^C in the wood formed during drought (Farquhar et al., 1989; Grossiord et al., 2014). Thus, drought stress can be quantified as increase in wood carbon isotope ratio (Δδ^13^C) between drought and normal years. Hence, growth responses and Δδ^13^C combined provide a powerful tool to quantify drought effects on trees.

Tree species vary greatly in their susceptibility to drought due to physiological and morphological differences. Among other features such as fine-root distribution and their die-back in response to drought (Brunner, Herzog, Dawes, Arend, & Sperisen, 2015; Sánchez-Pérez, Lucot, Bariac, & Trémolières, 2008), two key factors that might drive tree species reactions to drought are stomatal control and resistance to cavitation (Choat et al., 2012; Martínez-Vilalta & Garcia-Forner, 2017; McDowell et al., 2008). Stomatal closure in response to water deficits enables plants to avoid critically low water potentials through transpiration losses and thus hydraulic failure but species differ largely in their type of stomatal control (Martínez-Vilalta & Garcia-Forner, 2017; McDowell et al., 2008): Isohydric or water saving species close their stomata fast during water shortage, while anisohydric or water spending species keep their stomata open and continue to transpire (Martínez-Vilalta & Garcia-Forner, 2017). Next to stomatal control, xylem resistance to cavitation is a key determinant of tree responses to drought as embolism decreases water availability, which leads to desiccation and at extreme levels to tree death (Choat et al., 2012). It is conceivable that stomatal control and cavitation resistance interact, as a water spending behavior necessitates a continued water uptake via roots and, all else being equal, carries an increased risk for xylem cavitation (McDowell et al., 2008). However, whether this translates into water spending species exhibiting generally higher cavitation resistance and *vice versa* still remains elusive as some studies found indications for such a correlation (Klein, 2014; Martínez-Vilalta & Garcia-Forner, 2017) while others did not (Kröber, Zhang, Ehmig, & Bruelheide, 2014). We expect water saving species to show earlier growth and Δδ^13^C responses, while water spending species may face high cavitation risks during severe and prolonged drought conditions characterized by very low soil moisture availability. Hence, for understanding and generalizing the effects of consecutive droughts on forests, tree species should be examined that differ in such traits.

The high tree species richness of floodplain forests (Ward, Tockner, & Schiemer, 1999) makes them ideally suited for comparative studies of tree species reactions to consecutive droughts as they are one of the few systems where coexisting mature trees spanning an entire gradient of hydraulic behaviors can be found. Floodplain forests rank among the most rapidly disappearing ecosystems due to land conversion and drainage (Leuschner & Ellenberg, 2017; Mikac et al., 2018) and novel climatic conditions – like prolonged droughts – may amplify this trend through changing the hydrological regimes on which these forests depend. For instance, sinking groundwater levels may increase tree growth sensitivity to drought and susceptibility to drought-induced dieback (Mikac et al., 2018; Skiadaresis, Schwarz, & Bauhus, 2019) and this might bring these forests, which are among the most dynamic, productive and diverse Central European habitats (Kowalska et al., 2020; Tockner & Stanford, 2002), closer to a tipping point. On the contrary, the higher water availability in floodplain forests may buffer drought effects to a certain extent as trees might have access to groundwater in addition to precipitation-derived moisture (Heklau, Jetschke, Bruelheide, Seidler, & Haider, 2019). Hence, it is conceivable that if drought effects on growth and Δδ^13^C were observed in floodplain trees, other forest ecosystems might experience even stronger effects.

Here, we focus on the effect of the two consecutive drought years 2018–2019 characterized by extremely hot and dry conditions (Fig. 1), as well as their cumulative effects, on tree growth and Δδ^13^C as physiological stress response. To this end, we reconstruct the stress exerted by this unprecedented event and compare it to past (single) drought events based on tree-ring records from the dominant tree species – *Quercus robur*, *Acer pseudoplatanus* and *Fraxinus excelsior* – in the Leipzig floodplain forest, one of the few remaining and thus highly protected floodplain forests in Central Europe (BMU & BfN, 2021; Günther-Diringer et al., 2021). We sampled trees in two environmental strata representing topographic differences in distance to groundwater. We expect the results for the hypotheses proposed below to be more pronounced in the drier stratum. Specifically, we tested the following hypothesis:

**Fig. 1.**
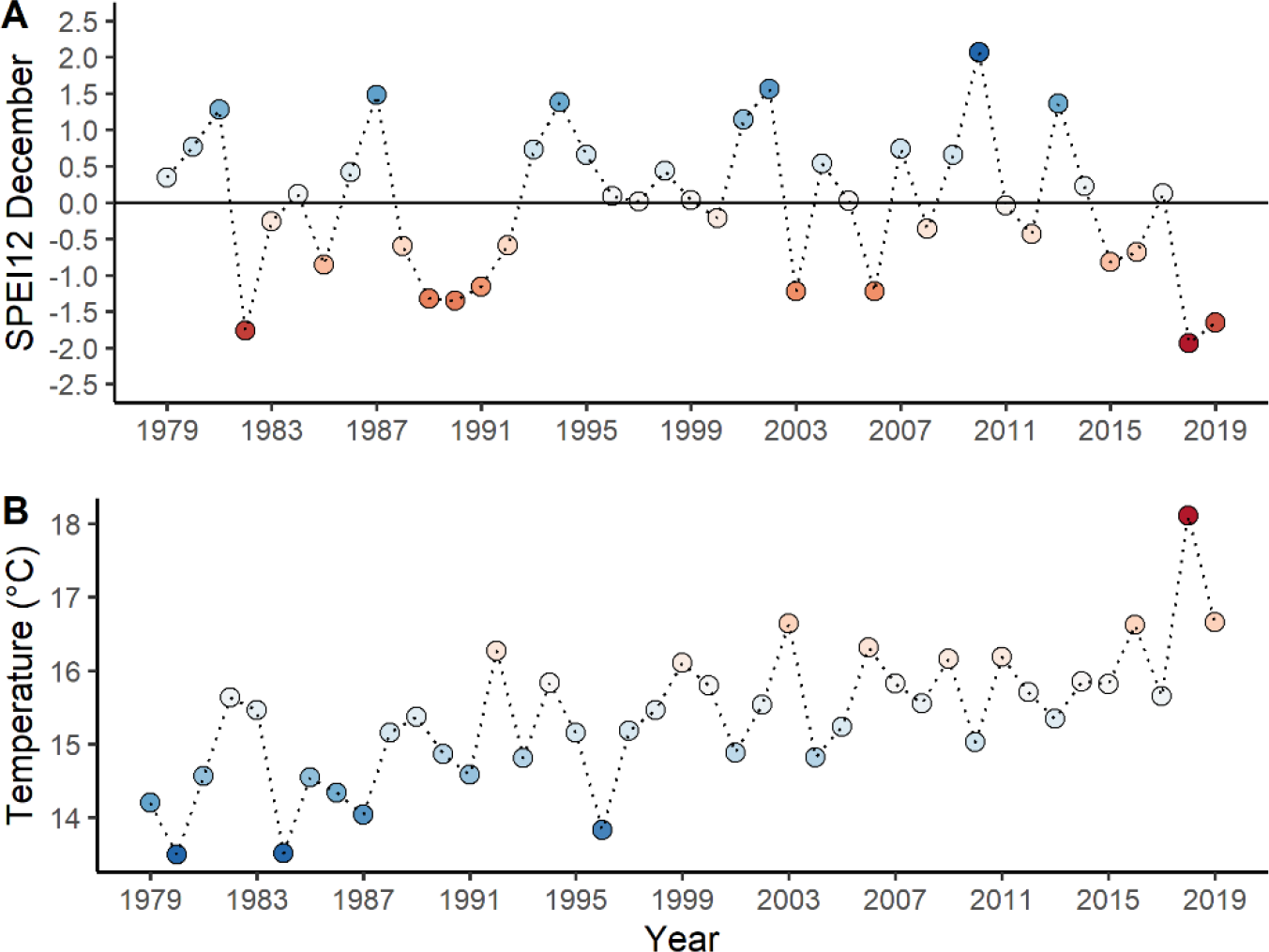
Annual standardized water balance of precipitation minus potential evapotranspiration (A, January-December) and mean growing season temperature (B, April-September) per year from 1979-2019 in the Leipzig floodplain forest. The water balance was calculated as Standardized Precipitation Evapotranspiration Index (SPEI; Vicente-Serrano, Beguería, & López-Moreno, 2010). Points are colored according to their value with deeper red indicating increasing drought and temperature. The horizontal line in (A) represents the long-term mean, negative values indicate water deficits and positive values water surpluses. SPEI values below -1 and above 1 can be considered exceptionally dry and wet, respectively. See Figs. S1–S2 for additional SPEI lengths, climatic and hydrological variables that we used to identify drought events.

**H1** Drought stress responses in 2018 – measured as growth reduction and a positive Δδ^13^C – are comparable to stress responses during former drought years.

**H2** The consecutive drought years 2018–2019 lead to a drastic growth reduction and a further increase in Δδ^13^C in 2019.

**H3** Water spending species respond faster to drought stress (already in 2018), while water saving species react later but show stronger reactions to the consecutive drought in 2019.

## Methods

### Study site

In this study, we used data collected from a Central European floodplain forest ecosystem located in the northwest of the city of Leipzig, Germany. The Leipzig floodplain forest is one of the few remaining and thus highly protected floodplain forests in Central Europe (BMU & BfN, 2021; Günther-Diringer et al., 2021) and lies in the transition zone between maritime and continental climate characterized by warm summers, with an annual mean temperature of 9.6 °C and an annual precipitation sum of 525 mm (1979–2019; DWD, Station Leipzig/Halle). Its main rivers Weiße Elster, Luppe, Pleiße and Parthe formed the floodplain landscape, but their course and thus the floodplain forest itself has been strongly influenced by human interventions over the last centuries (Gutte, 2011). The straightening of rivers as well as dike and canal constructions strongly influenced the hydrological regime of the floodplain forest, which today does not experience regular flooding anymore (Haase & Gläser, 2009). The floodplain soils originated from an accumulation of alluvial sediments, such as gravel, sand and loam, as result of several glacial periods (Haase & Gläser, 2009). These are nowadays covered by an alluvial clay layer with a thickness between 1–4 m, rich in nutrients and with a high pH (around 6–7) (Gutte, 2011; Haase & Gläser, 2009). The principal soil available to trees is thus a loamy Vega, with partly gleyed conditions, above gravel and sand filled with groundwater.

### Tree species

The contemporary floodplain forest ecosystem can be characterized as *Ficario-Ulmetum* Knapp ex Medwecka-Kornas 1952 with oak, elm and ash being the dominant tree species (Härdtle et al., 2020). The absence of flooding, however, resulted in an on-going gradual shift to an oak-hornbeam forest (*Galium-carpinetum stachyetosum*) and allowed other tree species (especially maple), which are intolerant to flooding, to become dominant. Moreover, elm (*Ulmus minor*) largely disappeared from the tree canopy due to the Dutch Elm Disease since the 1960s. Nowadays, the dominant tree species of Leipzig’s floodplain forest are *Quercus robur L.* (hereafter oak), *Acer pseudoplatanus L.* (hereafter maple) and *Fraxinus excelsior L.* (hereafter ash) (Haase & Gläser, 2009; Richter, Reu, Wirth, Doktor, & Vohland, 2016), on which we focus in the present study. These three species feature contrasting adaptations to drought in terms of stomatal control and cavitation resistance, which allowed us to explore a range of species response strategies to consecutive drought stress. In terms of stomatal control, former studies classified oak (Cocozza et al., 2020; Thomsen, Reisdorff, Gröngröft, Jensen, & Eschenbach, 2020) and maple (Köcher, Gebauer, Horna, & Leuschner, 2009; Lemoine, Peltier, & Marigo, 2001; Leuschner, Wedde, & Lübbe, 2019) as rather water saving (isohydric), while ash was shown to follow a water spending (anisohydric) strategy (Köcher et al., 2009; Lemoine et al., 2001; Leuschner et al., 2019). Consistent with this view, studies focussing on sap flow measurements on mature trees, which can be considered as proxy for tree transpiration and hence stomatal aperture, reported a significant downregulation of sap flow with decreasing soil water availability for maple but not for ash (Brinkmann, Eugster, Zweifel, Buchmann, & Kahmen, 2016; Hölscher, Koch, Korn, & Leuschner, 2005; Köcher et al., 2009), indicating water saving versus water spending modes, respectively. Importantly, we view stomatal control here as a gradient and not as a dichotomy between water saving or water spending behaviour (but see Martínez-Vilalta & Garcia-Forner, 2017), with classifications depending strongly on the compared species. However, quantitative data on traits indicative of stomatal control like sap flow is scarce (particularly for *Q. robur*) and we are not aware of any study that compared mature individuals of all herein examined species under similar site conditions. We therefore assembled high-resolution sap flow and soil moisture data recorded during the 2018 drought in the Leipzig floodplain forest (Leipzig Canopy Crane facility) to provide a quantitative comparison of species-specific sensitivity to decreasing soil moisture under severe drought (Fig. 2a–b) (Schnabel et al., 2021a). Consistent with the classification above, oak and maple significantly downregulated sap flow under drought conditions (indicating water saving behaviour) while ash maintained similar sap flow rates (indicating water spending behaviour) (Fig. 2b). In terms of species resistance to cavitation, we relied on published values of the water potential at which 50% of xylem conductivity is lost due to cavitation (Ψ50, Choat et al., 2012), the most common measure of embolism resistance in trees (Choat et al., 2012). This comparison indicates a similar cavitation resistance in oak (-2.8 MPa) and ash (-2.8 MPa) while maple is less resistant (-1.6 MPa).

**Fig. 2.**
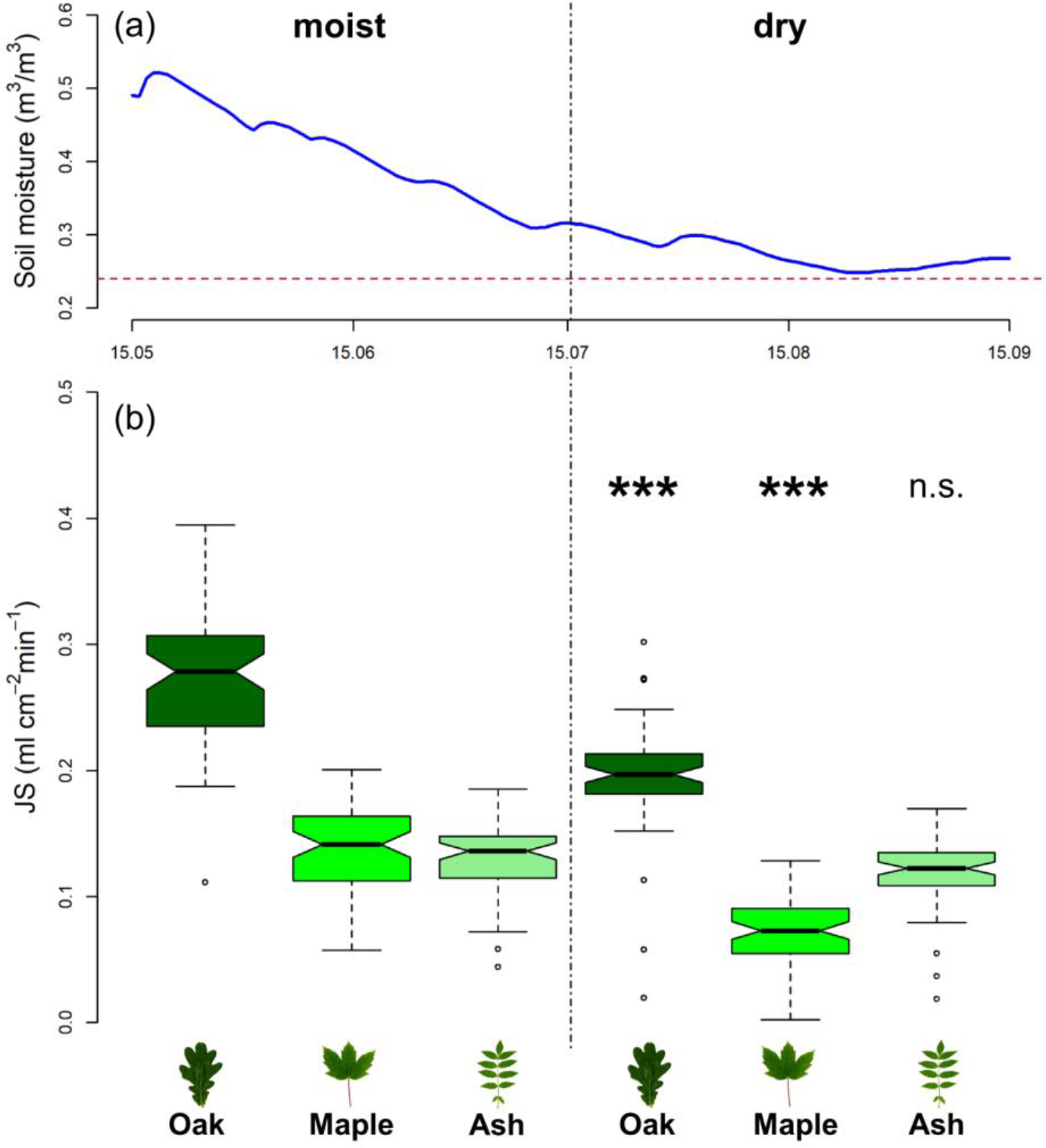
Soil moisture development (a) and sap flux density (J*s*) regulation of oak, maple and ash (b) during the 2018 hotter drought in the Leipzig floodplain forest. The observed soil moisture development (a) was used to delineate two periods with contrasting soil moisture conditions, a moist and dry period, respectively. During the dry period soil moisture levels approached 0.24 m^3^/m^3^ (red horizontal line), the permanent wilting point of vegetation on clay soils (Weil & Brady, 2017). Boxplots (b) show daily maxima in J*s* during the two month period with moist soil (mid-Mai to mid-July) and during the period with dry soil at later stages during the 2018 drought (mid-July to mid-September). A statistically significant downregulation of J*s* under dry compared to moist conditions is indicated by asterisks over the respective species’ boxplot (’***’ p <0.001; ’**’ p <0.01; ’* ’p <0.05). See Methods S1 for details on the sap flow and soil moisture measurements and analyses.

### Drought year identification

The definition and identification of drought is central to the analysis of drought effects. Here, we define drought as period with water deficits compared to normal conditions, where ‘normal’ can be quantified as a percentile of the long-term mean of meteorological or hydrological variables (Schwarz et al., 2020; van Loon et al., 2016). Following suggestions by Schwarz et al. (2020) we selected drought years based on climatic and hydrological information alone without considering tree growth reductions to avoid a biased selection that could for example result in the exclusion of drought years without reduced growth. We used the Standardized Precipitation Evapotranspiration Index (SPEI; Vicente-Serrano et al., 2010) and river discharge data to identify drought years. The SPEI is a commonly used drought index (Hari et al., 2020; Schwarz et al., 2020; Skiadaresis et al., 2019) based on the standardized monthly water balance of precipitation minus potential evapotranspiration. It can quantify drought severity according to a droughts intensity and duration and can be calculated at different time scales (e.g. 1–12 months) (Vicente-Serrano et al., 2010). Here, we used three different SPEI lengths that represent the climatic water balance of the main vegetation period (SPEI for 3 months, Mai–July), the full vegetation period (SPEI for 6 months, April–September) and the full year (SPEI for 12 months, January–December) for each year and with a 40-year reference period (1979–2019; Fig. S1). SPEI series were calculated with the SPEI package (Beguería & Vicente-Serrano, 2017) in R from monthly precipitation (mm) and potential evapotranspiration (mm) data derived from the weather station located closest to the study sites (DWD Climate Data Center [CDC], Station Leipzig/Halle, ID 2932; see Fig. S1 for details).

We classified years with SPEI values ≤ -1 as drought years, years with SPEI values ≥ 1 as particularly wet and years with values between -1 and 1 as ‘normal’ (McKee, Doesken, & Kleist, 1993). To take into account the hydrological regime of the floodplain forest, which is in addition to local precipitation strongly influenced by its rivers, we compared the SPEI derived classification to river discharge calculated for the same periods as the SPEIs (Fig. S2). We considered only years without particularly high discharge as drought years. Focusing on a 20-year period before the 2018–2019 consecutive drought, we selected 2005, 2009 and 2017 as reference years with normal climatic conditions, while single drought years – in contrast to the 2018–2019 consecutive drought

– were 2003, 2006 and 2015 (Fig. 1; Figs. S1–S2; hereafter ‘single drought years’). We did not consider a longer period to minimize the effect of past forest management and ground water fluctuation related influences on tree growth. Both, the drought in 2018 and the one in 2019 were the most severe droughts in the last 40 years (i.e. they had the lowest SPEI values), but 2018 had slightly lower SPEI values and was especially characterized by an extreme heat wave during the vegetation period.

### Tree selection and increment core extraction

We selected trees for extracting wood increment cores from permanent forest research plots of the “Lebendige Luppe” (living Luppe river) project (Scholz et al., 2018), which cover a gradient in topographic distances to the groundwater level (Fig. S3). The project features three distinct strata of distance to groundwater: dry (> 2m), intermediate (1–2m) and moist (≤ 1m) plots, with 20 plots per stratum each 0.25ha in size. Plots were not flooded since 1973 due to flood control measures (dikes, river-straightening etc.), except for winter 2011 and summer 2013, when the area of Leipzig experienced extreme flood events. We chose to sample trees on dry and moist plots to cover both ends of the gradient of hydrological site conditions within the floodplain forest (Fig. S3). Across these plots, we extracted tree increment cores from at least 40 tree individuals per species (20 trees per stratum) from each of the three dominant tree species oak, maple and ash, amounting to 120 sampled trees. From each tree, we extracted one increment core at a height of 80cm with a ∅ 5mm increment corer (Suunto, Sweden) in January–February 2020, i.e. in the winter after tree ring formation of the second consecutive drought year 2019 was completed. Trees with diameters at breast height (dbh) > 20cm were selected according to their dominance, past management history and health status. Competition for light is a central determinant of tree growth and δ^13^C that might complicate the detection of drought effects (Grossiord et al., 2014). We therefore sampled only dominant and co-dominant individuals, i.e. trees belonging to category 1–2 according to the classification of Kraft (1884), that were no direct competitors and further excluded plots that showed signs of forest management in recent years. We further selected only healthy appearing trees, excluding those ash trees visually affected by ‘ash dieback’ (*Hymenoscyphus fraxineus*) and those maple trees visually affected by the ‘sooty bark disease’ (*Cryptostroma corticale*). Both fungal pathogens had caused widespread tree damages and diebacks in the Leipzig floodplain forest during the 2018–2019 consecutive drought and especially very few ash trees were completely unaffected (Wirth et al., 2021). We used the classification key of Lenz, Straßer, Baumann, and Baie (2012) for ash dieback infestation and sampled only trees showing no to only little signs of infestation (level 0–2 of infestation levels 0–5) based on annual infestation records for four years prior to sampling. Importantly, our sample is thus representative for the most vital individuals of the entire population. Since the number of trees fulfilling these strict criteria was too low within the plot area, we sampled also oak and maple trees in the direct vicinity of the plots.

### Tree growth analysis

Tree cores were dried at 70 °C for at least 3 days and then clamped in wooden alignment strips. For surface preparation, we used a core microtome (WSL, Switzerland) (Gärtner & Nievergelt, 2010) to enhance visibility of tree ring boundaries. Tree ring-width was measured with a LINTAB 6 measuring table and the TSAPWin Professional 4.64 program © 2002–2009 Frank Rinn / RINNTECH with an accuracy of 1/1000 mm. The measured sequences were cross-dated against a species-specific master chronology developed in former works for the same area as well as against each other using COFECHA (Grissino-Mayer, 2001). This allowed us to identify missing rings, which were more often found in maple trees and in the consecutive drought years 2018–2019. Years without growth were included as zero for the respective year. Sequences that could not be dated unequivocally were excluded from further analysis. The final number of trees included for growth analysis was 114 trees, including 40 oak, 32 maple and 42 ash trees from 11 moist and 15 dry plots. Mean series length was 109 years for oak, 79 years for maple and 94 years for ash trees (Table S1).

Tree ring-width provides an integrated record of past growth conditions as influenced by environmental factors including but not limited to climate and shows an inherent decrease in ring-width with increasing tree size (Schweingruber, 1996). As we focus here on climatic influences on growth, we removed age-related trends from the raw tree ring-width chronologies via a Negative Exponential curve (Fritts, 1976), which provided the best compromise between removing long-term age trends and preserving decadal variability in growth using the package dplR (Bunn, 2008; Bunn et al., 2020). We assessed the climatic sensitivity of tree growth through computing bootstrapped Pearson’s correlation functions between species-specific chronologies and monthly climatic variables (Fig. S4), using the package treeclim (Zang & Biondi, 2015). Species-specific chronologies are shown in Fig. S5.

We quantified the growth response of trees to the consecutive drought years 2018 and 2019 as well as to single drought years (hereafter ‘growth response’) for each individual tree using the growth resistance index proposed by Lloret et al. (2011) as:

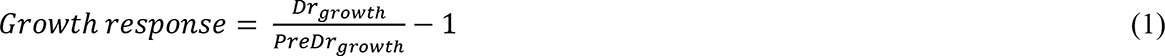

where *Drgrowth* is a tree’s detrended growth in drought year(s) and *PreDrgrowth* is a tree’s detrended growth in the reference period characterized by normal climatic conditions. The growth response index is thus standardized around zero with positive values meaning higher and negative values lower growth during drought year(s) compared to reference years. We calculated the growth response for 2018, 2019 and for single drought years and used the mean growth response in single drought years as baseline against which we compared the growth responses in 2018 and 2019. Recent decades experienced an unprecedented surge in temperatures and drought events (Buras et al., 2020; Schuldt et al., 2020), making the use of a continuous multiyear reference period before drought events that is not influenced by drought itself difficult. We therefore used the mean growth in three years (2005, 2009, 2017) that were characterized by normal climatic conditions and not proceeded by a drought year (see above) to calculate *PreDrgrowth*. We used several years to calculate *PreDrgrowth* and the growth response in single drought years to reduce the influence of outliers caused by individual tree reactions to factors other than climate (e.g. changes in competitive interactions, waterlogging). However, as the choice of growth data (i.e. using different detrending methods) and length of reference period can strongly influence results of the index used (Schwarz et al., 2020), we tested for the robustness of here reported relationships. We examined growth responses in detail based on raw and detrended ring-width and further compared growth responses calculated with a one-year pre-period (2017 was the only climatically normal year before the 2018–2019 consecutive drought; Fig. S1) to growth responses calculated with the mean reference period detailed above.

We quantified drought legacies in tree growth as observed growth minus predicted growth as expected based on the climatic water balance in the year after the drought event (Anderegg et al., 2015; Kannenberg et al., 2019). Tree growth in the last 40 years (1979–2019) was predicted using tree-specific regressions between detrended tree ring-width and SPEI12 of December (Fig. S1). We consistently used the twelve-month long SPEI of December to capture the climatic water balance of the full year (January–December) for each species to provide estimates of legacy effects that are comparable between species and drought years. Reported drought legacy effects thus quantify the deviation of observed growth from expected growth based on climate in year 1 after single drought years (2003, 2006, 2015) and after the hotter drought in 2018, that is, in the years 2004, 2007, 2016 (using their mean as baseline) and in the consecutive drought year 2019. Climatic conditions in year 1 after all single drought years were neither exceptionally dry nor wet (Fig. 1), providing a suitable baseline against which the legacy effects of consecutive drought can be compared. Finally, we compared our baseline modelling approach, which used the mean growth response and drought legacy effect in individual drought years, to an analysis that considered each single drought year individually.

### Carbon isotope analysis

The stable carbon isotope composition (δ¹³C) in wood of the same cores was measured following tree ring-width measurements. The tree rings of the herein analysed consecutive drought years 2018–2019, of single drought years (2003, 2006, 2015) and of reference years (2005, 2009, 2017) were separated and their wood tissue homogenized. Some individuals, especially maple trees, did not form tree rings during the 2018–2019 consecutive drought, likely due to intense drought stress. As their δ¹³C could thus not be analysed, we excluded these trees from our isotope analysis (6 maple and 1 oak tree). The homogenized material of the tree rings in reference years (2005, 2009 and 2017) and single drought years (2003, 2006 and 2015), was pooled by mixing equal shares of the material from each of the three years. The isotope analysis was done at the BGC stable isotope laboratory of the Max Planck Institute for Biogeochemistry in Jena, Germany. The results were expressed as isotopic ratio δ¹³C, calculated with the equation of Farquhar et al. (1989) as follows:

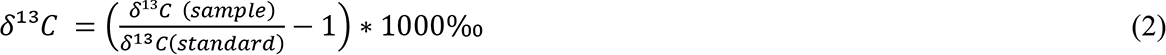

where *δ¹³C (sample)* and *δ¹³C (standard)* are the abundance ratios between ¹³C and ¹²C of the given sample and Vienna PeeDee Belemnite international standard (VPDB). Isotope ratios were expressed in δ-notation in per mil units (‰). We calculated the increase in δ¹³C from reference to drought years for each individual tree as indicator of a tree’s physiological stress response to drought as:

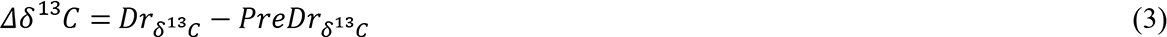

where *Drδ¹³C* is the isotope composition in drought year(s) and *PreDrδ¹³C* the isotope composition in the reference years (see e.g. Grossiord et al., 2014). Positive values of Δδ^13^C thus indicate higher and negative values lower stress during drought year(s) compared to reference years. Drought and reference years used to calculate Δδ¹³C were the same as in the growth response analysis.

### Statistical analysis

We used linear mixed-effects models (LMMs) to understand the effects of consecutive drought years on tree growth and Δδ¹³C in comparison to single drought years (using their mean growth response and Δδ¹³C in all analysis). We were further interested in understanding how these effects were modulated by changes in a trees distance to groundwater. We fitted species-specific LMMs for analysing the growth response and Δδ¹³C with the packages lme4 (Bates, Mächler, Bolker, & Walker, 2015) and lmerTest (Kuznetsova, Brockhoff, & Christensen, 2017) and a significance level of α=0.05. Drought event (single droughts, 2018, 2019), groundwater level (dry > 2m, moist ≤ 1m) and their interaction were modelled as fixed effects, while tree identities nested within plots were used as nested random effects to account for differences between plots and for repeated measurements on each individual tree. We selected the most parsimonious model structure via backward model selection, first adjusting the random (likelihood ratio tests) and then the fixed effect model structure (F-tests), using the step function in lmerTest. The most parsimonious LMM structure consistently retained only a fixed effect of drought event and tree identity nested within plot as random effect, indicating that water table did not significantly (P>0.05) influence observed relationships. Only for the growth response LMM of oak we found a significant interaction (P=0.0413) between drought event and groundwater level, which however disappeared when using non-detrended growth data or a one-year reference period. Therefore, to report only the most robust relationships, we present all final LMMs with drought event as the only fixed effect. Final LMMs (Table S2) were fit using restricted maximum likelihood estimation (REML) and marginal means and confidence intervals (95%) were predicted with the ggeffects package (Lüdecke, 2018). We used post-hoc pairwise comparisons with adjusted p-values for multiple comparisons (Tukey’s Honest Significant Difference) to compare differences between drought events using the emmeans package (Lenth, 2020) (Table S3). Model assumptions (normality, independence and homogeneity of variance) where visually checked through examining model residuals and through quantile-quantile plots. Drought legacy effects were analysed using the same modelling procedure (see Table S2). All analyses were conducted in R version 4.1.0 (R Core Team, 2021). Tree growth, δ¹³C and drought legacy data and analysis scripts are available via the iDiv data repository (Schnabel et al., 2021b).

## Results

We found pronounced responses to drought stress in terms of tree growth and Δδ^13^C across the examined tree species, with strongest stress responses in the second of two consecutive hotter drought years (2019). The mean growth response to single drought years (2003, 2006 and 2015) before the 2018–2019 consecutive drought ranged around zero for oak and maple, while growth in ash tended to be reduced (Fig. 3a–c). This indicates a similar tree growth in single drought years and in climatically ‘normal’ years for oak and maple but not for ash. Growth of oak and maple even tended to be higher in 2018 compared to normal years (mean growth response above zero). The hotter drought in 2018 did not induce growth responses in oak and maple that differed significantly from single drought years (P>0.1 for both species) but ash experienced an overall significantly stronger growth reduction (t=-2.94, P=0.004; Fig. 3c; Table S2). In 2019, the second consecutive and extreme drought year, the growth reduction in all species was significantly stronger than in single drought years (oak t=-2.00, P=0.049; maple t=-2.74, P=0.008; ash t=-7.22, P<0.001; Fig. 3a–c; Table S2) and in comparison to 2018 (Table S3). Observed growth responses were largely insensitive to the type of growth data (raw or detrended) and reference period (one-year or pooled years) used (Figs. S6–S7). We used species-specific models but provide evidence for significant differences between species in Fig. S8. Distance to groundwater had an overall small influence on the growth response of the examined species (non-significant effect of groundwater level for maple and ash). Only for oak we found indications for a smaller growth response on moist plots in 2019 (significant interaction of drought year and groundwater level, P=0.041). Of the three analysed species, ash, followed by maple, showed a high growth sensitivity to drought (especially to SPEI series of summer months indicating summer drought) while oak was the least sensitive (Fig. S4). Moreover, high summer temperatures negatively affected the growth of ash and maple but not of oak (Fig. S4).

**Fig. 3.**
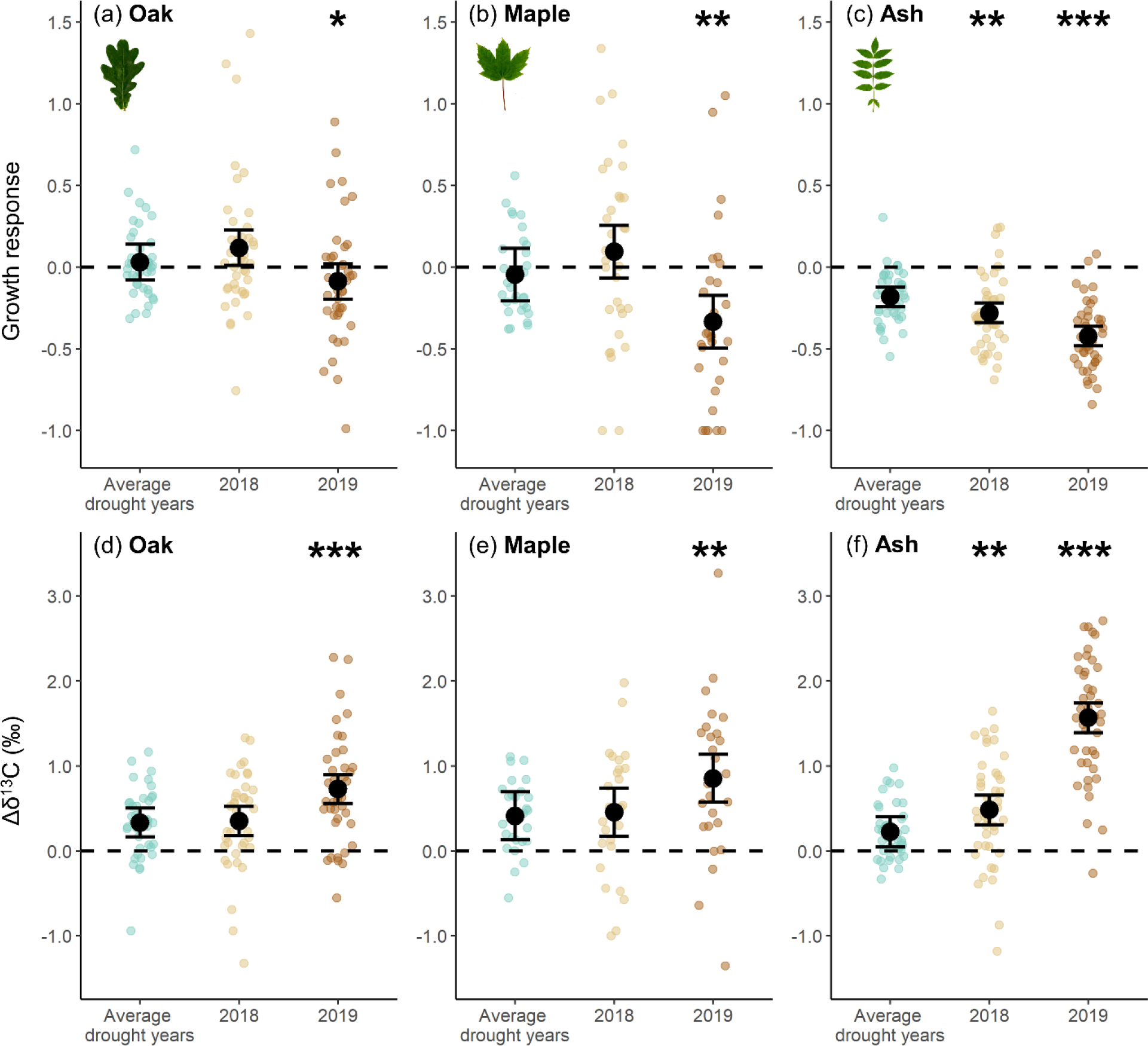
Growth response and increase in the carbon isotope ratio (Δδ^13^C) in wood of oak, maple and ash in drought years. The figure shows the growth response (upper panels) and Δδ^13^C (lower panels) in the consecutive hotter drought years 2018 and 2019 compared to the mean growth response and Δδ^13^C in single drought years (2003, 2006 and 2015). Zero corresponds to a comparable growth and δ^13^C in drought and climatically normal years. Negative growth response values indicate growth reductions while positive Δδ^13^C values indicate stress increases during drought compared to normal years. The growth response and Δδ^13^C were calculated with Eq. 1 and Eq. 3, respectively. Black points show estimated marginal means and error bars the 95% confidence intervals of linear mixed-effects model fits, with non-overlapping confidence intervals indicating signficant differences. Coloured points show the growth response and Δδ^13^C values per tree and species (oak n=40, n=39; maple n=32, n=26; ash n=42, n=42) and are jittered to enhance visibility. The tree ring-widths have been detrended with a negative exponential function. Statistically significant differences in the growth response and Δδ^13^C between the years 2018 and 2019 compared to single drought years are indicated by asterisks over the respective year (’***’ p <0.001; ’**’ p <0.01; ’* ’p <0.05).

We did not find drought legacy effects in tree growth after single drought years, that is, observed tree growth in year 1 after these droughts was not significantly lower than growth predicted based on climate (Fig. 4a–c; Table S2). For ash, observed growth even tended to be higher than predicted (Fig. 4c). In contrast, the hotter drought year 2018 induced substantial legacy effects in maple and ash but not in oak (Fig. 4a–c), that is, observed growth in the second consecutive drought year 2019 was significantly lower than expected based on climate. Legacy effects in 2018 were significantly different from legacy effects in single drought years for maple and ash but not for oak (oak t=-0.93, P=0.358; maple t=-3.52, P<0.001; ash t=-8.41, P<0.001; Fig. 4a–c; Table S2). Here presented models that used the mean across single drought years (baseline models; Figs. 3–4; Table S3) yielded similar conclusions as models that analysed each single drought year individually (Figs. S9–10; Table S4).

**Fig. 4.**
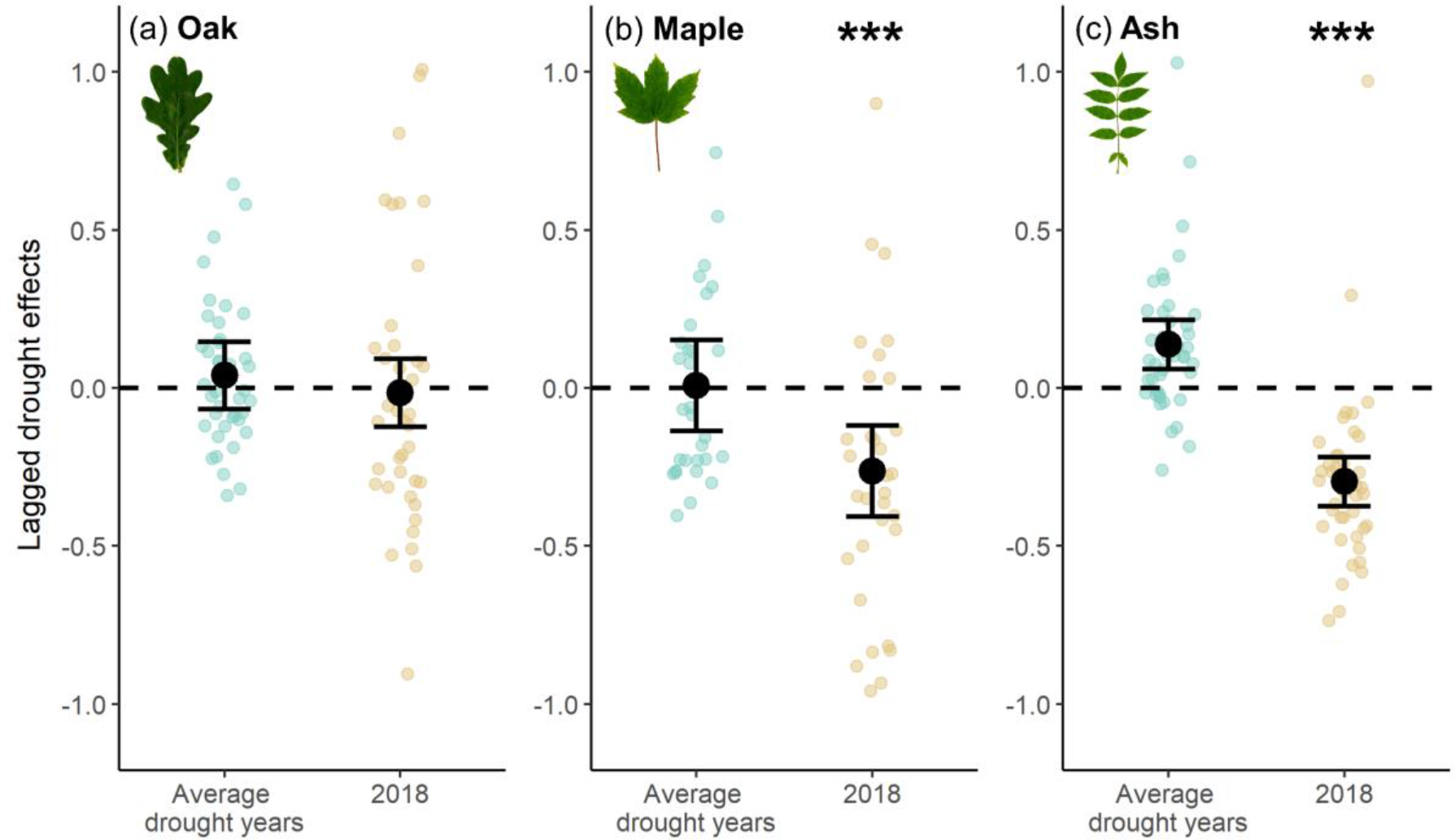
Drought legacy effects in growth of oak, maple and ash in the year following drought events. The figure shows drought legacy effects induced by the hotter drought year 2018 compared to mean legacy effects induced by single drought years (2003, 2006 and 2015). Legacy effects were quantified as observed minus predicted (detrended) tree ring-width based on climate in year 1 after the drought event. Zero corresponds to growth as expected based on climate conditions, while negative values indicate drought legacies in form of lower than expected post-drought growth. Black points show estimated marginal means and error bars the 95% confidence intervals of linear mixed-effects model fits, with non-overlapping confidence intervals indicating signficant differences. Coloured points show legacy effects per tree and species (oak n=40; maple n=32; ash n=42) and are jittered to enhance visibility. Statistically significant differences in legacy effects between 2018 compared to single drought years are indicated by asterisks (’***’ p <0.001; ’**’ p <0.01; ’* ’p <0.05).

We found positive Δδ^13^C values across all species and drought years, indicating tree physiological stress responses to drought irrespective of drought type (single or consecutive; Fig. 3d–f). However, the magnitude of Δδ^13^C increases varied strongly between drought years and species. For oak and maple, Δδ^13^C was not significantly enhanced in 2018 compared to single drought years (P=0.85 and P=0.79), while ash had significantly higher Δδ^13^C values (t=2.85, P=0.006; Fig. 3f; Table S2). Across all species, we found a strong increase in Δδ^13^C in 2019 compared to single drought years (oak t=3.93, P<0.001; maple t=2.80, P=0.007; ash t=14.80, P<0.001; Fig. 3d–f; Table S2) and in comparison to 2018 (Table S3). The Δδ^13^C increase was strongest for ash. Distance to groundwater had no significant influence on Δδ^13^C for all examined species. Together these results indicate that drought stress in 2018 was, except for ash, comparable to stress in former single drought years, while the second consecutive drought year 2019 induced the strongest growth reductions and increases in Δδ^13^C across all species.

## Discussion

Using tree growth reductions and increases in Δδ^13^C as indicators of drought stress, we report a strong increase in drought-related stress in the second of two consecutive hotter drought years across all examined species. Drought responses were consistent for both indicators (growth response and Δδ^13^C; Fig. 3), but the timing and magnitude of responses were species-specific: Oak showed the overall smallest stress response followed by maple with the strongest response in ash. The 2019 drought, although an extreme drought as well, was meteorologically less severe than the preceding drought year 2018 (Fig. 1). This and observed drought legacy effects (Fig. 4) indicate that the cumulative drought effect exerted by both years was likely the principal driver of the stress increase in 2019. The 2018 hotter drought was the severest drought so far recorded in Central Europe (Buras et al., 2020; Hari et al., 2020; Schuldt et al., 2020), but, as predicted, we found physiological stress increases (Δδ^13^C) to be comparable to former single drought years and tree growth to be largely within the range of climatically ‘normal’ years. Hence, the comparably high water availability in floodplain forests may partly buffer tree stress responses to single but not to consecutive drought years.

Our conclusion that the effects of single drought years were buffered to some extent contrasts with the dramatic drought effects reported across European forests in 2018 that suffered widespread defoliation, xylem hydraulic failure and mortality (Buras et al., 2020; Schuldt et al., 2020) but is consistent with other floodplain forest studies. For instance, the exceptionally high gross primary production during the warm spring in 2018 was found to compensate for losses later that year due to drought in a Czech floodplain forest (Kowalska et al., 2020). Similarly, tree growth recovered within two years after the 1976 drought for all herein analyzed tree species, which was attributed to the buffering effect of water availability in floodplain forests (Heklau et al., 2019). Nonetheless, we found physiological stress increases (Δδ¹³C) in 2018 while tree growth in most species did not react. This confirms the view of clearer drought signals in Δδ¹³C compared to tree ring-width, potentially due to tree growth being maintained from carbon reserves even under low soil water availability (Jucker et al., 2017).

This picture changed dramatically in 2019. As hypothesized, we observed the strongest stress responses in the second consecutive drought year. Drought legacy effects (Anderegg et al., 2015) were found to be widespread in forests and to affect tree growth and Δδ¹³C 1–5 years after the actual drought event (Anderegg et al., 2013; Anderegg et al., 2015; Gazol et al., 2020; Kannenberg et al., 2019; Lloret et al., 2011; Szejner et al., 2020). We observed significant drought legacy effects in tree growth after the 2018 hotter drought but not after former single drought years. Hence, in a system where drought legacy effects have not been observed previously, the hotter drought in 2018 was severe enough to induce such legacies. It should be noted that the reference drought years (2003, 2006, 2015) were themselves considered as some of the severest droughts in Central Europe (Allen et al., 2015; Büntgen et al., 2021). Their comparably low effect thus supports our view of high water availability in floodplain forests partly buffering tree stress responses and simultaneously underlines the unprecedented nature of 2018–2019. Former studies on drought legacy effects examined post drought periods in which trees where already (partially) recovering (e.g. Gazol et al., 2020). In contrast, we focus here on two consecutive hotter drought years, unprecedented in severity for at least since 250 years (Hari et al., 2020), which left the trees no time to recover. The few studies that studied prolonged droughts, moreover, did not examine the cumulative built-up of drought effects from year-to-year as they used either mean tree growth across drought years or growth in the last year of drought to calculate growth responses to drought (Schwarz et al., 2020). In comparison, the strong reactions we report for 2019 should be mainly attributable to legacy effects of 2018 (see also some early reports of drought legacies in Buras et al. (2020) and Schuldt et al. (2020)). Other changes in the trees’ environment like reduced competition for light are unlikely within a single year. In addition, forest management can be excluded as potential cause as we did not sample trees in stands that experienced recent interventions.

Several physiological mechanisms could explain drought legacy effects (Anderegg et al., 2015) and thereby cumulative drought stress. Drought-induced xylem cavitation may impair growth and transpiration (and thus effect Δδ¹³C) (McDowell et al., 2008; Schuldt et al., 2020). Under consecutive drought, this damage persists, while vulnerability to cavitation may continue to increase under successive drought stress (Anderegg et al., 2013). In the second drought year, less nonstructural carbohydrates (NSC) reserves were likely left for xylem repair, growth and especially for keeping up the trees’ defense system, which increases their susceptibility to pests and pathogens (Anderegg et al., 2013; Hartmann & Trumbore, 2016; McDowell et al., 2008; Schuldt et al., 2020). Although we studied only the most vital tree individuals of the population, thus largely excluding disease effects from our sample, the majority of ash trees in the forest were affected to some degree (Wirth et al., 2021). It is therefore not possible, to completely disentangle whether the species intrinsic traits, incipient ash-dieback or their interaction caused the strong stress response in this species. Drought induces shifts in carbon allocation in favor of the canopy and root system at the expense of radial growth (Brunner et al., 2015; Kannenberg et al., 2019), for instance to replace fine roots lost during drought (Brunner et al., 2015), which would reduce tree-ring growth and thereby amplify drought legacy effects. Finally, when photosynthesis is insufficient to meet demands, NSC reserves are utilized to maintain autotrophic respiration, growth and tissue repair (Hartmann & Trumbore, 2016; Richardson et al., 2013). This enriches the reserve pool and tissues built from it in ^13^C as the isotopically lighter ^12^C is turned over faster than ^13^C, which may have further contributed to the strong increase in Δδ¹³C in 2019 in addition to fractionation through stomata closure.

In addition to physiological drought legacies, different meteorological and abiotic conditions may have contributed to the strong stress responses observed in 2019. The second hotter drought year 2019 started already with severe soil moisture shortages as the low winter and spring precipitation in 2018–2019 was not enough to refill soil water reservoirs (UFZ Drought Monitor / Helmholtz Centre for Environmental Research). The drought legacy effects we found in tree growth therefore likely resulted from both, physiological and abiotic drought legacies. Next to drought duration and intensity, drought timing may influence tree radial growth (Schwarz et al., 2020). We observed variable timings of climatic drought onset, with single drought years being characterized by both spring and summer droughts, 2018 by summer drought (onset in May) and 2019 by drought during spring and summer (onset in February) (Fig. S11). Studies examining intra-annual radial growth at high temporal resolution show that maple, ash and oak species continue to grow until August (if not affected by drought) (Brinkmann et al., 2016; Dietrich, Delzon, Hoch, Kahmen, & Battipaglia, 2018), which, together with our own observation of strong growth-climate correlations in spring and summer months (Fig. S4), points at all species being effected by drought during their growing phase. Nonetheless, drought effects on growth and Δδ¹³C are likely strongest if the timing of drought is such that both early and latewood development are affected (Schwarz et al., 2020). That the drought in 2019 affected the entire growing season while the drought in 2018 did not, may therefore — in addition to legacy effects of 2018 — have contributed to the strong stress responses we report. Our sampling sites cover the whole gradient of groundwater conditions in the examined floodplain forest but interestingly we found only small effects of groundwater level. The reasons remain speculative. Differences in distances to the groundwater level may have been too small to induce strong effects on tree performance or, alternatively, more intense rooting on dry plots may have compensated for lower water availability (Skiadaresis et al., 2019). We did not observe a temporal trend in groundwater levels (neither decrease nor increase) over the study period and decreases in response to the 2018–2019 drought were small (Fig. S12). Temporal changes in groundwater level are thus unlikely to have had major influences on observed responses. Finally, trees in floodplain forests take up water from upper (unsaturated) soil horizons that are fed by precipitation and through capillary rise from the groundwater level (Sánchez-Pérez et al., 2008; Singer et al., 2014). Under severe drought this capillary rise may have been interrupted at all distances from the groundwater while the loamy Vega soils of our study site reached moisture levels close to the permanent wilting point of vegetation for clay soils (Fig. 2a).

The magnitude and timing of drought stress responses were species-specific, which may be related to differences in species hydraulic traits. Oak and ash feature similar cavitation resistance but different stomatal control which may explain the stronger drought stress response observed in ash compared to oak. We report a highly significant downregulation of sap flow with decreasing soil moisture during the 2018 drought in oak and maple but no downregulation in ash (Fig. 2), indicating rather water saving and water spending strategies, respectively. Its water saving strategy may have helped oak to avoid xylem cavitation during peak drought periods, while the water spending strategy of ash carried a higher cavitation risk (Martínez-Vilalta & Garcia-Forner, 2017; McDowell et al., 2008), as both species feature similar levels of cavitation resistance (both -2.8 MPa; Choat et al., 2012). Moreover, a water spending strategy necessitates continued water uptake via roots (McDowell et al., 2008), which may be an especially risky strategy on severely dried out clay soils.

Oak and maple showed similar Δδ¹³C responses in all drought years consistent with their similar stomatal control. In contrast, ash showed a stronger response particularly in 2019. On first sight, this may come as a surprise as one may expect lower Δδ¹³C increases (which are related to stomatal closure) in a water spending compared to water saving species. However, potentially high hydraulic damages in ash during the severe 2018 drought would necessitate a high mobilization of NSC reserves for damage repair. As discussed above, this would enrich the reserve pool and tissues built from it in Δδ¹³C and could explain the strong Δδ¹³C increases we observe in the second consecutive drought year 2019. Future studies should directly measure NSC dynamics during drought to confirm these expectations. The overall intermediate drought reaction of maple, which is often considered drought sensitive (Leuschner et al., 2019), may be related to its higher vulnerability to cavitation and/or its water saving behavior that may have prevented severe damages to a certain extent. Moreover, the reaction of maple may also be influenced by its less exposed crown position (maple trees were rather co-dominant) which can reduce irradiance and water pressure deficits (Montgomery, Reich, & Palik, 2010). Finally, short-term growth responses to drought need to be contextualized. We found growth of ash and maple to be sensitive to both features of hotter droughts (low water availability and high temperatures) while oak was insensitive to either factor during the last 40 years (Fig. S4). However, this does not mean that oak does not react to drought, but rather that its response is non-linear as highlighted by its unprecedented response to the 2019 drought.

Other traits may have influenced the responses observed but establishing species-specific differences remains challenging. For instance, ash was reported to have fine- and coarse-root biomass concentrated to shallower soil layers than oak in another riparian hardwood forests (Sánchez-Pérez et al., 2008). However, other studies reported rather deep rooting in ash and an intruding ability to plastically shift its water uptake to deeper soil layers (Brinkmann, Eugster, Buchmann, & Kahmen, 2019; Meißner, Köhler, Schwendenmann, & Hölscher, 2012). Similarly, fine-root die-back is, just as aboveground leaf shedding, a common tree response to drought (Brunner et al., 2015; Kuster, Arend, Günthardt-Goerg, & Schulin, 2013; Meier & Leuschner, 2008). It thus likely contributed to herein reported drought responses but we lack data on species- and site-specific differences to test this hypothesis. Despite compelling progress in functional trait research (Kattge et al., 2020), assessments of key drought tolerance traits, particularly fine root and stomatal control related ones, thus remain scarce and should be a research priority in future studies including at our study sites. Finally, here reported drought effects may be influenced through the naturally high tree species richness of floodplain forests (Ward et al., 1999), as diverse tree communities with dissimilar hydraulic traits may outperform species poor communities through complementarity in water use (Sánchez-Pérez et al., 2008; Schnabel et al., 2019).

## Conclusion

The response of forests to the increasing frequency and intensity of droughts (IPCC, 2014) will affect a variety of ecosystem services and will determine if forests act as carbon sink or source in the 21^st^ century. Our retrospective analysis based on tree rings allowed us a robust comparison of the cumulative stress responses observed in the hotter drought years 2018–2019 compared to responses in former severe drought years (2003, 2006, 2015) on the same tree individuals. Tree stress responses in 2019 were stronger than in any other examined drought year, indicating that consecutive hotter drought years exert a novel stress. Comparisons of living and dead trees affected by drought show that radial growth reductions are widespread before tree mortality and that sudden changes in tree growth often precede mortality caused by tree hydraulic failure (Cailleret et al., 2017; Obladen et al., 2021). Against this background it is important to consider that we found partly buffered tree stress responses, presumably because floodplain trees are fed by groundwater in addition to precipitation, and examined only the most vital tree individuals of the population. Our results thus show a ‘best-case scenario’ and more severe tree responses, such as widespread tree mortality, could be expected if entire tree populations or other forest ecosystems were examined (see e.g. Buras et al. (2020), Schuldt et al. (2020), Wirth et al. (2021)). Further, it remains unknown how the here observed responses will affect tree recovery after and resilience to (future) drought, but the reported persistence of legacy effects for years (Anderegg et al., 2015) is worrying. Nonetheless, a species like oak that combines a high tolerance to drought and flood (Scharnweber, Manthey, & Wilmking, 2013), may remain resilient, underlining its importance for floodplain forests. Consecutive hotter droughts are projected to become more frequent (Hari et al., 2020). Results of this and similar research may contribute towards forecasting tree species and forest responses to this novel climatic phenomenon.

## Supporting information

Supplementary material for Schnabel et al.

## Acknowledgements

We thank our colleagues of the ‘Lebendige Luppe’ (living Luppe river) project for establishing the plot network that allowed us to draw representative conclusions and Andreas Sickert (Stadtforst) und Andreas Padberg (Sachsenforst) for the possibility to extract tree increment cores. The Lebendige Luppe project is funded by the German Federal Agency for Nature Conservation (BfN) in cooperation with the German Federal Ministry for the Environment, Nature Conservation and Nuclear Safety (BMU). We further acknowledge the help of the BGC stable isotope laboratory of the Max Planck Institute for Biogeochemistry in Jena, Germany. F.S. was supported by the International Research Training Group TreeDì funded by the Deutsche Forschungsgemeinschaft (DFG, German Research Foundation) – 319936945/GRK2324.

## Author contributions

C.W., F.S., R.A.E., A.K., R.R., C.S.-D. and G.S. conceived the idea and developed the concept of the study. S.P., L.S., A.K., F.S., R.A.E., R.R. and C.S.-D. collected data. F.S., S.P, L.S., R.R. and G.S. analyzed data. F.S., C.S.-D. and G.S. created figures. F.S. wrote the manuscript with contributions from S.P. and L.S. All authors discussed the results and contributed substantially to revisions.

## Data availability

Data supporting the findings in this study is archived in public repositories. Tree growth, δ¹³C and drought legacy data and analysis scripts (Schnabel et al., 2021b) are publicly available via the iDiv Data Repository: https://idata.idiv.de/ddm/Data/ShowData/3497?version=0. Soil moisture and sap flux data and analysis scripts (Schnabel et al., 2021a) are available upon reasonable request via the iDiv Data Repository: https://idata.idiv.de/ddm/Data/ShowData/3498?version=0. Climate data is publicly available at CDC (Station Leipzig/Halle, ID 2932): https://cdc.dwd.de/portal/.

## References

Allen, C. D., Breshears, D. D., & McDowell, N. G. (2015). On underestimation of global vulnerability to tree mortality and forest die-off from hotter drought in the Anthropocene. Ecosphere, 6(8). https://doi.org/10.1890/ES15-00203.1

Allen, C. D., Macalady, A. K., Chenchouni, H., Bachelet, D., McDowell, N., Vennetier, M., … Cobb, N. (2010). A global overview of drought and heat-induced tree mortality reveals emerging climate change risks for forests. Forest Ecology and Management, 259(4), 660–684. https://doi.org/10.1016/j.foreco.2009.09.001

Anderegg, W. R. L., Plavcová, L., Anderegg, L. D. L., Hacke, U. G., Berry, J. A., & Field, C. B. (2013). Drought’s legacy: Multiyear hydraulic deterioration underlies widespread aspen forest die-off and portends increased future risk. Global Change Biology, 19(4), 1188–1196. https://doi.org/10.1111/gcb.12100

Anderegg, W. R. L., Schwalm, C., Biondi, F., Camarero, J. J., Koch, G., Litvak, M., … Pacala, S. (2015). Pervasive drought legacies in forest ecosystems and their implications for carbon cycle models. Science, 349(6247), 528–532.

Anderegg, W. R. L., Trugman, A. T., Badgley, G., Anderson, C. M., Bartuska, A., Ciais, P., … Randerson, J. T. (2020). Climate-driven risks to the climate mitigation potential of forests. Science, 368(6497). https://doi.org/10.1126/science.aaz7005

Bates, D., Mächler, M., Bolker, B., & Walker, S. (2015). Fitting Linear Mixed-Effects Models Using lme4. Journal of Statistical Software, 67(1). https://doi.org/10.18637/jss.v067.i01

Beguería, S., & Vicente-Serrano, S. M. (2017). SPEI: Calculation of the Standardised Precipitation-Evapotranspiration Index. [Computer software]. Retrieved from https://CRAN.R-project.org/package=SPEI

BMU & BfN (2021). Auenzustandsbericht. Flussauen in Deutschland. Berlin, Germany.

Brinkmann, N., Eugster, W., Buchmann, N., & Kahmen, A. (2019). Species-specific differences in water uptake depth of mature temperate trees vary with water availability in the soil. *Plant Biology (Stuttgart*, Germany), 21(1), 71–81. https://doi.org/10.1111/plb.12907

Brinkmann, N., Eugster, W., Zweifel, R., Buchmann, N., & Kahmen, A. (2016). Temperate tree species show identical response in tree water deficit but different sensitivities in sap flow to summer soil drying. Tree Physiology, 36(12), 1508–1519. https://doi.org/10.1093/treephys/tpw062

Brunner, I., Herzog, C., Dawes, M. A., Arend, M., & Sperisen, C. (2015). How tree roots respond to drought. Frontiers in Plant Science, 6, 547. https://doi.org/10.3389/fpls.2015.00547

Bunn, A. G. (2008). A dendrochronology program library in R (dplR). Dendrochronologia, 26(2), 115–124. https://doi.org/10.1016/j.dendro.2008.01.002

Bunn, A. G., Korpela, M., Biondi, F., Campelo, F., Mérian, P., Qeadan, F., & Zang, C. (2020). dplR: Dendrochronology Program Library in R. Retrieved from https://CRAN.R-project.org/package=dplR

Büntgen, U., Urban, O., Krusic, P. J., Rybníček, M., Kolář, T., Kyncl, T., … Trnka, M. (2021). Recent European drought extremes beyond Common Era background variability. Nature Geoscience. Advance online publication. https://doi.org/10.1038/s41561-021-00698-0

Buras, A., Rammig, A., & Zang, C. S. (2020). Quantifying impacts of the 2018 drought on European ecosystems in comparison to 2003. Biogeosciences, 17(6), 1655–1672. https://doi.org/10.5194/bg-17-1655-2020

Cailleret, M., Jansen, S., Robert, E. M. R., DeSoto, L., Aakala, T., Antos, J. A., … Martínez-Vilalta, J. (2017). A synthesis of radial growth patterns preceding tree mortality. Global Change Biology, 23(4), 1675–1690. https://doi.org/10.1111/gcb.13535

Choat, B., Jansen, S., Brodribb, T. J., Cochard, H., Delzon, S., Bhaskar, R., … Zanne, A. E. (2012). Global convergence in the vulnerability of forests to drought. Nature, 491(7426), 752–755. https://doi.org/10.1038/nature11688

Cocozza, C., Paoletti, E., Mrak, T., Zavadlav, S., Levanič, T., Kraigher, H., … Hoshika, Y. (2020). Isotopic and Water Relation Responses to Ozone and Water Stress in Seedlings of Three Oak Species with Different Adaptation Strategies. Forests, 11(8), 864. https://doi.org/10.3390/f11080864

Dietrich, L., Delzon, S., Hoch, G., Kahmen, A., & Battipaglia, G. (2018). No role for xylem embolism or carbohydrate shortage in temperate trees during the severe 2015 drought. Journal of Ecology, 197(4), 1142. https://doi.org/10.1111/1365-2745.13051

DWD Climate Data Center (CDC). Monthly mean of station observations of precipitation in mm, air temperature at 2 m above ground in °C, daily air temperature minimum at 2 m above ground in °C, daily air temperature maximum at 2 m above ground in °C, wind force in bft, cloud coverage in eighth, air pressure at station level in hPa, relative humidity at 2 m above ground in %, vapor pressure in hPa for Germany, version v19.3, last accessed: 06.02.2020. Data for Station Leipzig/Halle, ID 2932.

Farquhar, G. D., Ehleringer, J. R., & Hubick, K. T. (1989). Carbon Isotope Discrimination and Photosynthesis. Annual Review of Plant Physiology and Plant Molecular Biology, 40(1), 503– 537. https://doi.org/10.1146/annurev.pp.40.060189.002443

Fritts, H. (1976). Tree Rings and Climate. London, UK: Academic Press. Retrieved from http://site.ebrary.com/lib/alltitles/docDetail.action?docID=10685739

Gärtner, H., & Nievergelt, D. (2010). The core-microtome: A new tool for surface preparation on cores and time series analysis of varying cell parameters. Dendrochronologia, 28(2), 85–92. https://doi.org/10.1016/j.dendro.2009.09.002

Gazol, A., Camarero, J. J., Sangüesa-Barreda, G., Serra-Maluquer, X., Sánchez-Salguero, R., Coll, L., & Casals, P. (2020). Tree Species Are Differently Impacted by Cumulative Drought Stress and Present Higher Growth Synchrony in Dry Places. Frontiers in Forests and Global Change, 3. https://doi.org/10.3389/ffgc.2020.573346

Griscom, B. W., Adams, J., Ellis, P. W., Houghton, R. A., Lomax, G., Miteva, D. A., … Fargione, J. (2017). Natural climate solutions. Proceedings of the National Academy of Sciences of the United States of America, 114(44), 11645–11650. https://doi.org/10.1073/pnas.1710465114

Grissino-Mayer, H. D. (2001). Evaluating Crossdating Accuracy: A Manual and Tutorial for the Computer Program COFECHA. Tree-Ring Research. Retrieved from http://hdl.handle.net/10150/251654

Grossiord, C., Granier, A., Ratcliffe, S., Bouriaud, O., Bruelheide, H., Checko, E., … Gessler, A. (2014). Tree diversity does not always improve resistance of forest ecosystems to drought. Proceedings of the National Academy of Sciences of the United States of America, 111(41), 14812–14815. https://doi.org/10.1073/pnas.1411970111

Günther-Diringer, D., Berner, K., Koenzen, U., Kurth, A., Modrak, P., Ackermann, W., … Heyden, J. (2021). Methodische Grundlagen zum Auenzustandsbericht 2021: Erfassung, Bilanzierung und Bewertung von Flussauen. *BfN-Skripten*, 591.

Gutte, P. (2011). Das Querco-Ulmetum minoris ISSLER 1942, der Stieleichen-Ulmen- Hartholzwald, in der Elster-Luppe-Aue bei Leipzig. Mauritiana, 22, 213–242.

Haase, D., & Gläser, J. (2009). Determinants of floodplain forest development illustrated by the example of the floodplain forest in the District of Leipzig. Forest Ecology and Management, 258(5), 887–894. https://doi.org/10.1016/j.foreco.2009.03.025

Härdtle, W., Bergmeier, E., Fichtner, A., Heinken, T., Hölzel, N., Remy, D., … Dierschke, H. (2020). Pflanzengesellschaft des Jahres 2021: Hartholz-Auenwald (Ficario-Ulmetum). Tuexenia, 40, 373–399.

Hari, V., Rakovec, O., Markonis, Y., Hanel, M., & Kumar, R. (2020). Increased future occurrences of the exceptional 2018-2019 Central European drought under global warming. Scientific Reports, 10(1), 12207. https://doi.org/10.1038/s41598-020-68872-9

Hartmann, H., & Trumbore, S. (2016). Understanding the roles of nonstructural carbohydrates in forest trees - from what we can measure to what we want to know. New Phytologist, 211(2), 386–403. https://doi.org/10.1111/nph.13955

Heklau, H., Jetschke, G., Bruelheide, H., Seidler, G., & Haider, S. (2019). Species-specific responses of wood growth to flooding and climate in floodplain forests in Central Germany. IForest - Biogeosciences and Forestry, 12(3), 226–236. https://doi.org/10.3832/ifor2845-012

Hölscher, D., Koch, O., Korn, S., & Leuschner, C. (2005). Sap flux of five co-occurring tree species in a temperate broad-leaved forest during seasonal soil drought. Trees, 19(6), 628– 637. https://doi.org/10.1007/s00468-005-0426-3

IPCC (2014). Climate change 2014. Impacts, Adaptation, and Vulnerability. Part A: Global and Sectoral Aspects. Contribution of Working Group II to the Fifth Assessment Report of the Intergovernmental Panel on Climate Change. New York, NY: Cambridge University Press.

Jucker, T., Grossiord, C., Bonal, D., Bouriaud, O., Gessler, A., & Coomes, D. A. (2017). Detecting the fingerprint of drought across Europe’s forests: do carbon isotope ratios and stem growth rates tell similar stories? Forest Ecosystems, 4(1), 706. https://doi.org/10.1186/s40663-017-0111-1

Kannenberg, S. A., Novick, K. A., Alexander, M. R., Maxwell, J. T., Moore, D. J. P., Phillips, R. P., & Anderegg, W. R. L. (2019). Linking drought legacy effects across scales: From leaves to tree rings to ecosystems. Global Change Biology, 25(9), 2978–2992. https://doi.org/10.1111/gcb.14710

Kattge, J., Bönisch, G., Díaz, S., Lavorel, S., Prentice, I. C., Leadley, P., … Wirth, C. (2020). Try plant trait database - enhanced coverage and open access. Global Change Biology, 26(1), 119–188. https://doi.org/10.1111/gcb.14904

Klein, T. (2014). The variability of stomatal sensitivity to leaf water potential across tree species indicates a continuum between isohydric and anisohydric behaviours. Functional Ecology, 28(6), 1313–1320. https://doi.org/10.1111/1365-2435.12289

Köcher, P., Gebauer, T., Horna, V., & Leuschner, C. (2009). Leaf water status and stem xylem flux in relation to soil drought in five temperate broad-leaved tree species with contrasting water use strategies. Annals of Forest Science, 66(1), 101. https://doi.org/10.1051/forest/2008076

Kowalska, N., Šigut, L., Stojanović, M., Fischer, M., Kyselova, I., & Pavelka, M. (2020). Analysis of floodplain forest sensitivity to drought. Philosophical Transactions of the Royal Society of London. Series B, Biological Sciences, 375(1810), 20190518. https://doi.org/10.1098/rstb.2019.0518

Kraft, G. (1884). Beiträge zur Lehre von den Durchforstungen, Schlagstellungen und Lichtungshieben. Hannover, Germany: Klindworth’s Verlag.

Kröber, W., Zhang, S., Ehmig, M., & Bruelheide, H. (2014). Linking xylem hydraulic conductivity and vulnerability to the leaf economics spectrum--a cross-species study of 39 evergreen and deciduous broadleaved subtropical tree species. PLoS ONE, 9(11), e109211. https://doi.org/10.1371/journal.pone.0109211

Kuster, T. M., Arend, M., Günthardt-Goerg, M. S., & Schulin, R. (2013). Root growth of different oak provenances in two soils under drought stress and air warming conditions. Plant and Soil, 369(1-2), 61–71. https://doi.org/10.1007/s11104-012-1541-8

Kuznetsova, A., Brockhoff, P. B., & Christensen, R. H. B. (2017). lmerTest Package: Tests in Linear Mixed Effects Models. Journal of Statistical Software, 82(13). https://doi.org/10.18637/jss.v082.i13

Lemoine, D., Peltier, J. P., & Marigo, G. (2001). Comparative studies of the water relations and the hydraulic characteristics in Fraxinus excelsior, Acer pseudoplatanus and A. opalus trees under soil water contrasted conditions. Annals of Forest Science, 58(7), 723–731. https://doi.org/10.1051/forest:2001159

Lenth, R. (2020). emmeans: Estimated Marginal Means, aka Least-Squares Means. [Computer software]. Retrieved from https://CRAN.R-project.org/package=emmeans

Lenz, H., Straßer, L., Baumann, M., & Baie, U. (2012). Boniturschlüssel zur Einstufung der Vitalität von Alteschen. AFZ-DerWald, 3, 18–19.

Leuschner, C., & Ellenberg, H. (2017). *Vegetation ecology of Central Europe: Vegetation Ecology of Central Europe, Volume I* (L. Sutcliffe, Trans.). *Life sciences*. Cham, Switzerland: Springer.

Leuschner, C., Wedde, P., & Lübbe, T. (2019). The relation between pressure–volume curve traits and stomatal regulation of water potential in five temperate broadleaf tree species. Annals of Forest Science, 76(2). https://doi.org/10.1007/s13595-019-0838-7

Lloret, F., Keeling, E. G., & Sala, A. (2011). Components of tree resilience: Effects of successive low-growth episodes in old ponderosa pine forests. Oikos, 120(12), 1909–1920. https://doi.org/10.1111/j.1600-0706.2011.19372.x

Lüdecke, D. (2018). ggeffects: Tidy Data Frames of Marginal Effects from Regression Models. Journal of Open Source Software, 3(26), 772. https://doi.org/10.21105/joss.00772

Martínez-Vilalta, J., & Garcia-Forner, N. (2017). Water potential regulation, stomatal behaviour and hydraulic transport under drought: Deconstructing the iso/anisohydric concept. Plant, Cell and Environment, 40(6), 962–976. https://doi.org/10.1111/pce.12846

McDowell, N., Pockman, W. T., Allen, C. D., Breshears, D. D., Cobb, N., Kolb, T., … Yepez, E. A. (2008). Mechanisms of plant survival and mortality during drought: Why do some plants survive while others succumb to drought? New Phytologist, 178(4), 719–739. https://doi.org/10.1111/j.1469-8137.2008.02436.x

McKee, T. B., Doesken, N. J., & Kleist, J. (1993). The relationship of drought frequency and duration to time scales. In Proceedings of the 8th Conference on Applied Climatology.

Meier, I. C., & Leuschner, C. (2008). Belowground drought response of European beech: fine root biomass and carbon partitioning in 14 mature stands across a precipitation gradient. Global Change Biology, 14(9), 2081–2095. https://doi.org/10.1111/j.1365-2486.2008.01634.x

Meißner, M., Köhler, M., Schwendenmann, L., & Hölscher, D. (2012). Partitioning of soil water among canopy trees during a soil desiccation period in a temperate mixed forest. Biogeosciences, 9(8), 3465–3474. https://doi.org/10.5194/bg-9-3465-2012

Mikac, S., Žmegač, A., Trlin, D., Paulić, V., Oršanić, M., & Anić, I. (2018). Drought-induced shift in tree response to climate in floodplain forests of Southeastern Europe. Scientific Reports, 8(1), 16495. https://doi.org/10.1038/s41598-018-34875-w

Montgomery, R. A., Reich, P. B., & Palik, B. J. (2010). Untangling positive and negative biotic interactions: Views from above and below ground in a forest ecosystem. Ecology, 91(12), 3641–3655. https://doi.org/10.1890/09-1663.1

Obladen, N., Dechering, P., Skiadaresis, G., Tegel, W., Keßler, J., Höllerl, S., … Seim, A. (2021). Tree mortality of European beech and Norway spruce induced by 2018-2019 hot droughts in central Germany. Agricultural and Forest Meteorology, 307, 108482. https://doi.org/10.1016/j.agrformet.2021.108482

R Core Team (2021). A language and environment for statistical computing [Computer software]. Vienna, Austria: R Foundation for Statistical Computing: R Foundation for Statistical Computing. Retrieved from https://www.R-project.org/

Richardson, A. D., Carbone, M. S., Keenan, T. F., Czimczik, C. I., Hollinger, D. Y., Murakami, P., … Xu, X. (2013). Seasonal dynamics and age of stemwood nonstructural carbohydrates in temperate forest trees. New Phytologist, 197(3), 850–861. https://doi.org/10.1111/nph.12042

Richter, R., Reu, B., Wirth, C., Doktor, D., & Vohland, M. (2016). The use of airborne hyperspectral data for tree species classification in a species-rich Central European forest area. International Journal of Applied Earth Observation and Geoinformation, 52, 464–474. https://doi.org/10.1016/j.jag.2016.07.018

Sánchez-Pérez, J. M., Lucot, E., Bariac, T., & Trémolières, M. (2008). Water uptake by trees in a riparian hardwood forest (Rhine floodplain, France). Hydrological Processes, 22(3), 366–375. https://doi.org/10.1002/hyp.6604

Scharnweber, T., Manthey, M., & Wilmking, M. (2013). Differential radial growth patterns between beech (Fagus sylvatica L.) and oak (Quercus robur L.) on periodically waterlogged soils. Tree Physiology, 33(4), 425–437. https://doi.org/10.1093/treephys/tpt020

Schnabel, F., Purrucker, S., Schmitt, L., Engelmann, R. A., Kahl, A., Richter, R., … Wirth, C. (2021a). Cumulative growth and stress responses to the 2018-2019 drought in a European floodplain forest: Data and code on soil moisture and sap flux responses to drought. Retrieved from https://idata.idiv.de/ddm/Data/ShowData/3498?version=0

Schnabel, F., Purrucker, S., Schmitt, L., Engelmann, R. A., Kahl, A., Richter, R., … Wirth, C. (2021b). Cumulative growth and stress responses to the 2018–2019 drought in a European floodplain forest: Data and code on tree growth and δ13C responses to drought. Retrieved from https://idata.idiv.de/ddm/Data/ShowData/3497?version=0

Schnabel, F., Schwarz, J. A., Dănescu, A., Fichtner, A., Nock, C. A., Bauhus, J., & Potvin, C. (2019). Drivers of productivity and its temporal stability in a tropical tree diversity experiment. Global Change Biology, 25(12), 4257–4272. https://doi.org/10.1111/gcb.14792

Scholz, M., Seele, C., Engelmann, R. A., Hartmann, H., Heinrich, J. Henle, K., Herkelrath, A., … Riedel, J. (2018). Das Projekt Lebendige Luppe – ein Beitrag zur Renaturierung der Leipziger Nord-West-Aue. Auenmagazin, 14(14), 14–21.

Schuldt, B., Buras, A., Arend, M., Vitasse, Y., Beierkuhnlein, C., Damm, A., … Kahmen, A. (2020). A first assessment of the impact of the extreme 2018 summer drought on Central European forests. Basic and Applied Ecology, 45, 86–103. https://doi.org/10.1016/j.baae.2020.04.003

Schwarz, J. A., Skiadaresis, G., Kohler, M., Kunz, J., Schnabel, F., Vitali, V., & Bauhus, J. (2020). Quantifying Growth Responses of Trees to Drought—a Critique of Commonly Used Resilience Indices and Recommendations for Future Studies. Current Forestry Reports, 6, 185–200. https://doi.org/10.1007/s40725-020-00119-2

Schweingruber, F. H. (1996). Tree rings and environment: dendroecology. Berne, Switzerland: Paul Haupt AG Bern.

Singer, M. B., Sargeant, C. I., Piégay, H., Riquier, J., Wilson, R. J. S., & Evans, C. M. (2014). Floodplain ecohydrology: Climatic, anthropogenic, and local physical controls on partitioning of water sources to riparian trees. Water Resources Research, 50(5), 4490–4513. https://doi.org/10.1002/2014WR015581

Skiadaresis, G., Schwarz, J. A., & Bauhus, J. (2019). Groundwater Extraction in Floodplain Forests Reduces Radial Growth and Increases Summer Drought Sensitivity of Pedunculate Oak Trees (Quercus robur L.). Frontiers in Forests and Global Change, 2, 267. https://doi.org/10.3389/ffgc.2019.00005

Szejner, P., Belmecheri, S., Ehleringer, J. R., & Monson, R. K. (2020). Recent increases in drought frequency cause observed multi-year drought legacies in the tree rings of semi-arid forests. Oecologia, 192(1), 241–259. https://doi.org/10.1007/s00442-019-04550-6

Thomsen, S., Reisdorff, C., Gröngröft, A., Jensen, K., & Eschenbach, A. (2020). “Responsiveness of mature oak trees (Quercus robur L.) to soil water dynamics and meteorological constraints in urban environments”. Urban Ecosystems, 23(1), 173–186. https://doi.org/10.1007/s11252-019-00908-z

Tockner, K., & Stanford, J. A. (2002). Riverine flood plains: present state and future trends. Environmental Conservation, 29(3), 308–330. https://doi.org/10.1017/S037689290200022X

UFZ Drought Monitor / Helmholtz Centre for Environmental Research. Retrieved from https://www.ufz.de/index.php?en=37937

Van Loon, A. F., Stahl, K., Di Baldassarre, G., Clark, J., Rangecroft, S., Wanders, N., … van Lanen, H. A. J. (2016). Drought in a human-modified world: reframing drought definitions, understanding, and analysis approaches. Hydrology and Earth System Sciences, 20(9), 3631– 3650. https://doi.org/10.5194/hess-20-3631-2016

Vicente-Serrano, S. M., Beguería, S., & López-Moreno, J. I. (2010). A Multiscalar Drought Index Sensitive to Global Warming: The Standardized Precipitation Evapotranspiration Index. Journal of Climate, 23(7), 1696–1718. https://doi.org/10.1175/2009JCLI2909.1

Ward, J. V., Tockner, K., & Schiemer, F. (1999). Biodiversity of floodplain river ecosystems: ecotones and connectivity1. Regulated Rivers: Research & Management, 15(1-3), 125–139. https://doi.org/10.1002/(SICI)1099-1646(199901/06)15:1/3<125::AID-RRR523>3.0.CO;2-E

Weil, R. R., & Brady, N. C. (2017). Nature and properties of soils (15th edition). Pearson.

Wirth, C., Engelmann, R. A., Haack, N., Hartmann, H., Richter, R., Schnabel, F., … Seele-Dilbat, C. (2021). Naturschutz und Klimawandel im Leipziger Auwald: Ein Biodiversitätshotspot an der Belastungsgrenze. Biologie in Unserer Zeit, 51(1), 55–65. https://doi.org/10.11576/biuz-4107

Zang, C., & Biondi, F. (2015). treeclim: an R package for the numerical calibration of proxy- climate relationships. Ecography, 38(4), 431–436. https://doi.org/10.1111/ecog.01335

