## Supplementary material for Schnabel et al. for "Cumulative growth and stress responses to the 2018–2019 drought in a European floodplain forest"

**This file includes:**

Method S1

Figs. S1 to S12

Tables S1 to S4

References

**Method S1**

Soil moisture, that is, the volumetric water content, was measured at a depth of 0.1m in 10-minute intervals using a ML3 Theta-soil moisture sensor (Delta-T Devices Ltd, England) at the site of the Leipzig Canopy Crane facility in the Leipzig floodplain forest. The sensor was placed at a distance >3m from the nearest tree to minimize effects of stem flow. The observed soil moisture development (a) was used to delineate two periods with contrasting soil moisture conditions, a moist (mid-Mai to mid-July) and dry period (mid-July to mid-September), respectively. The latter period was characterized by soil moisture levels close to 0.24 $m^3/m^3$, the permanent wilting point of vegetation on clay soils (Weil & Brady, 2017).

Sap flux density (J*s*), which we consider as proxy for tree transpiration, was measured with heat dissipation sensors (UP Sap Flow-System, SFS2 TypM, UP Umweltanalytische Produkte GmbH, Germany) for three replicated tree individuals of oak (*Quercus robur*), maple (*Acer pseudoplatanus*) and ash (*Fraxinus excelsior*) at the site of the Leipzig Canopy Crane facility. At the north-facing side of trees (at 3.5m height) thermal probes were inserted to a depth of 2.5cm into the water-conducting sapwood. Sensors were protected from direct solar irradiation by reflective foil. Differences in temperature ($\Delta$T) between the heated (constant 12V current for power supply) and the unheated sensor were recorded at 10-minute intervals. $J_s$ (ml $cm^{-2}$ $min^{-1}$) was calculated from temperature differences according to (Granier, 1987) and species-specific daily maxima in $J_s$ were quantified as 95% quantile of $J_s$ values during each day. We tested for significant differences in $J_s$ between species and soil moisture period (moist and dry conditions) with analysis of variance. We found a highly significant interaction ($P<0.0001$) between species identity and soil moisture period. Significant differences in $J_s$ between the moist and dry soil moisture period for each species (asterisks lower panel Fig. 2) are based on post-hoc tests, with p-values adjusted for multiple comparisons using Tukey's Honest Significant Difference and a significance level of $\alpha$=0.05.

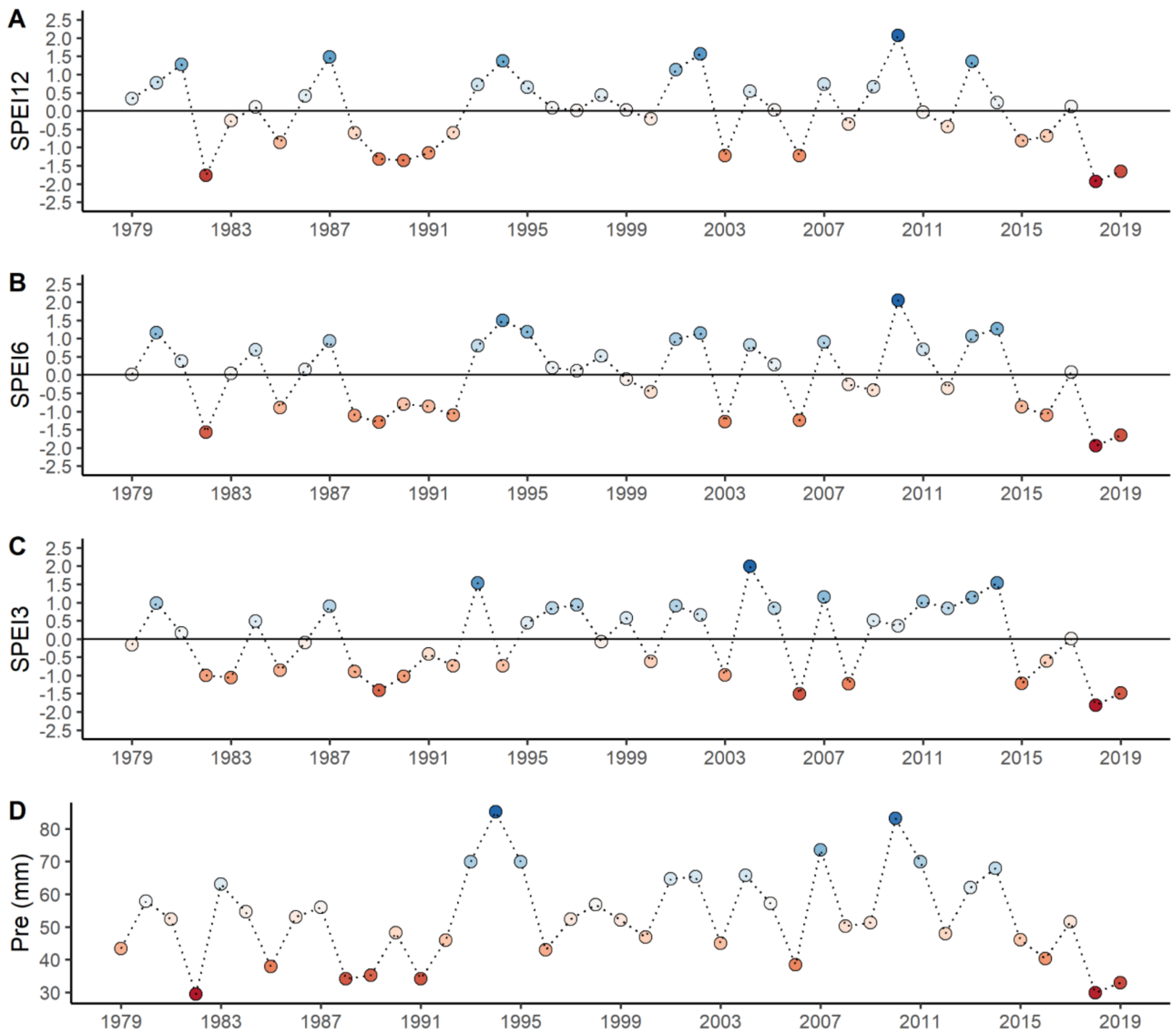


**Fig. S1** Standardized water balances of precipitation minus potential evapotranspiration (A-C, Standardized Precipitation Evapotranspiration Index (SPEI)) and mean monthly precipitation during the growing season (D, April-September) per year from 1979–2019 in the Leipzig floodplain forest. Points are colored according to their value with deeper red indicating increasing drought severity. Shown are SPEI values for each year at three time scales: (A) 12 months from January–December (SPEI12; see Fig. 1), (B) 6 months from April–September (SPEI6) and (C) 3 months from May–July (SPEI3). The horizontal line in (A–C) represents the long-term mean, negative values indicate water deficits and positive values water surpluses. SPEI values below -1 and above 1 can be considered as exceptionally dry and wet, respectively (McKee, Doesken, & Kleist, 1993). Potential evapotranspiration, which was not directly available from the DWD

Station Leipzig/Halle (ID 2932), was calculated with the FAO-56 Penman-Monteith equation (Beguería & Vicente-Serrano, 2017) using the following input data: monthly means of daily minimum temperature (°C), daily maximum temperature (°C), wind speed ($m\ s^{-1}$), cloud cover (%), air pressure (kPa), relative humidity (%), vapor pressure (kPa) as well as station elevation and latitude. All climate data  were provided by DWD Climate Data Center [CDC] and, where necessary, converted to the units described above.

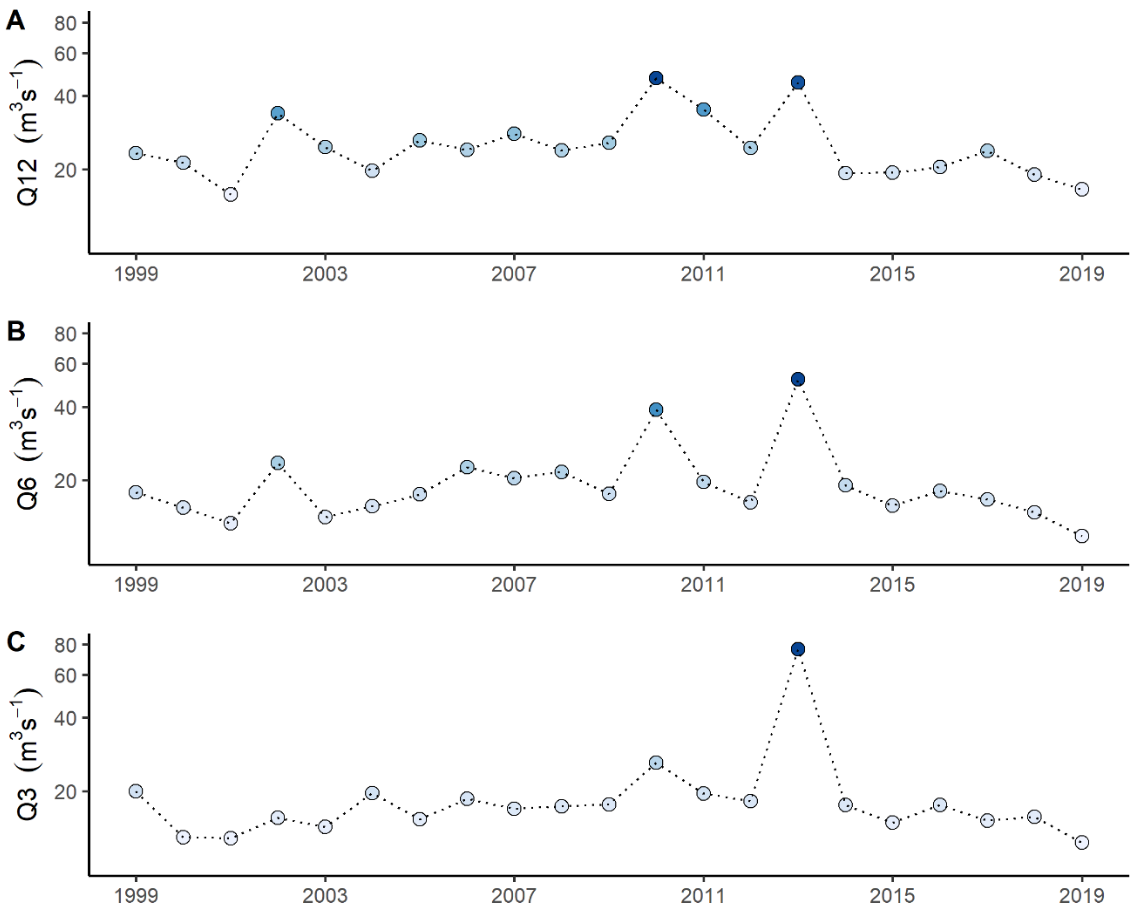


**Fig. S2** River discharge per year during the study period in the Leipzig floodplain forest. Shown is the mean daily discharge (log-scale) per year for the same three time scales used to characterize drought conditions (SPEIs; Supplementary Fig. 1): (A) 12 months from January–December (Q12), (B) 6 months from April–September (Q6) and (C) 3 months from May–July (Q3). Points are colored according to their value with deeper blue indicating increasing discharge. Note that the floodplain forest experienced an extreme flood event in 2013. River discharge was measured at the nearest official gauging station (Oberthau, LHW-576900) that is representative for the studied floodplain system.

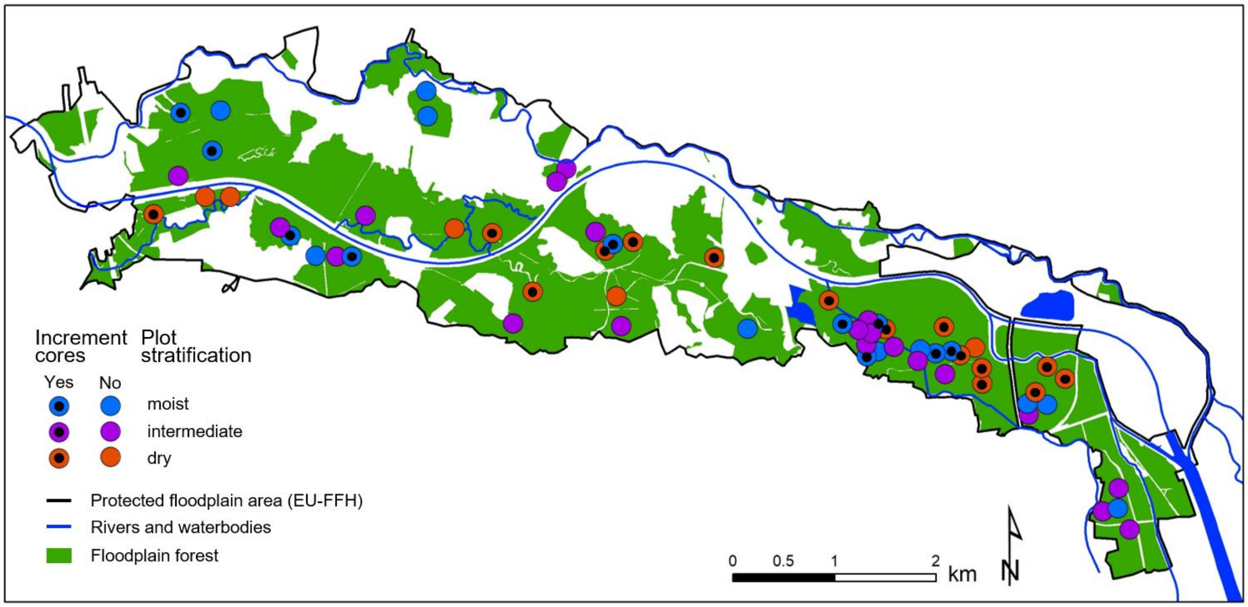


**Fig. S3** Map of study area within the Leipzig floodplain forest. Plots of the "Lebendige Luppe" (living Luppe river; Scholz et al., 2018) project are shown as dots, with different colours representing the plots distance to groundwater: dry (red, > 2m), intermediate (purple, 1–2m) and moist (blue, ≤ 1m). Plots from which tree increment cores were sampled are highlighted with black dots. Plots are located within the protected European Flora-Fauna Habitat floodplain area (EU-FFH 4639-301).

**Table S1** Mean length of tree ring chronologies in years ± one standard deviation for each species and stratum (dry, groundwater level > 2m; moist, groundwater level ≤ 1m).

| Species | Stratum | | |
|---|---|---|---|
| | moist | dry | mean |
| Oak | 102.9 ± 33.5 | 115.7 ± 29.8 | **109.3 ± 31.6** |
| Maple | 74.4 ± 23.1 | 83.1 ± 24.4 | **78.8 ± 23.8** |
| Ash | 95.9 ± 35.7 | 91.1 ± 26.1 | **93.5 ± 30.9** |

Note: Bold values show the mean value per species. The series length in years is an indicator of but not the same as the age of a tree. It corresponds only to the number of years that were visible on the extracted core.

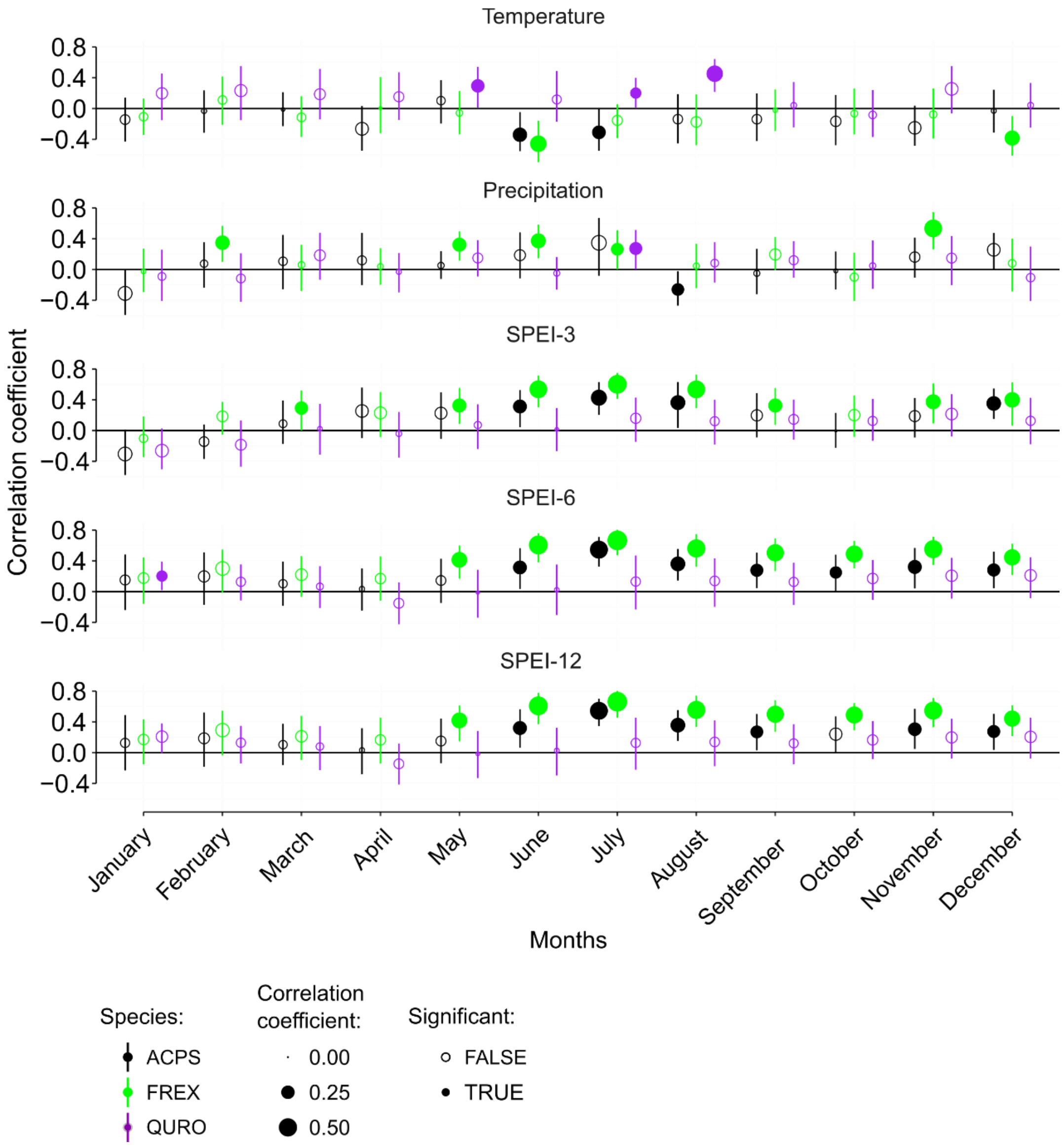


**Fig. S4** Climatic sensitivity of tree ring-width chronologies. Shown are bootstrapped Pearson's correlation coefficients between detrended species-specific ring-width chronologies and climate variables (temperature, precipitation, SPEI-3, SPEI-6 and SPEI-12) for the three species oak (QURO), maple (ACPS) and ash (FREX) for each month (January–December). Bars denote upper and lower confidence intervals (95%). Tree ring-width was detrended with a negative exponential curve. Detrending, chronology development and quality control were done with the dplR package (Bunn, 2008, 2010; Bunn et al., 2020) and correlation functions were computed over the period between 1979 and 2019 using the package treeclim (Zang & Biondi, 2015).

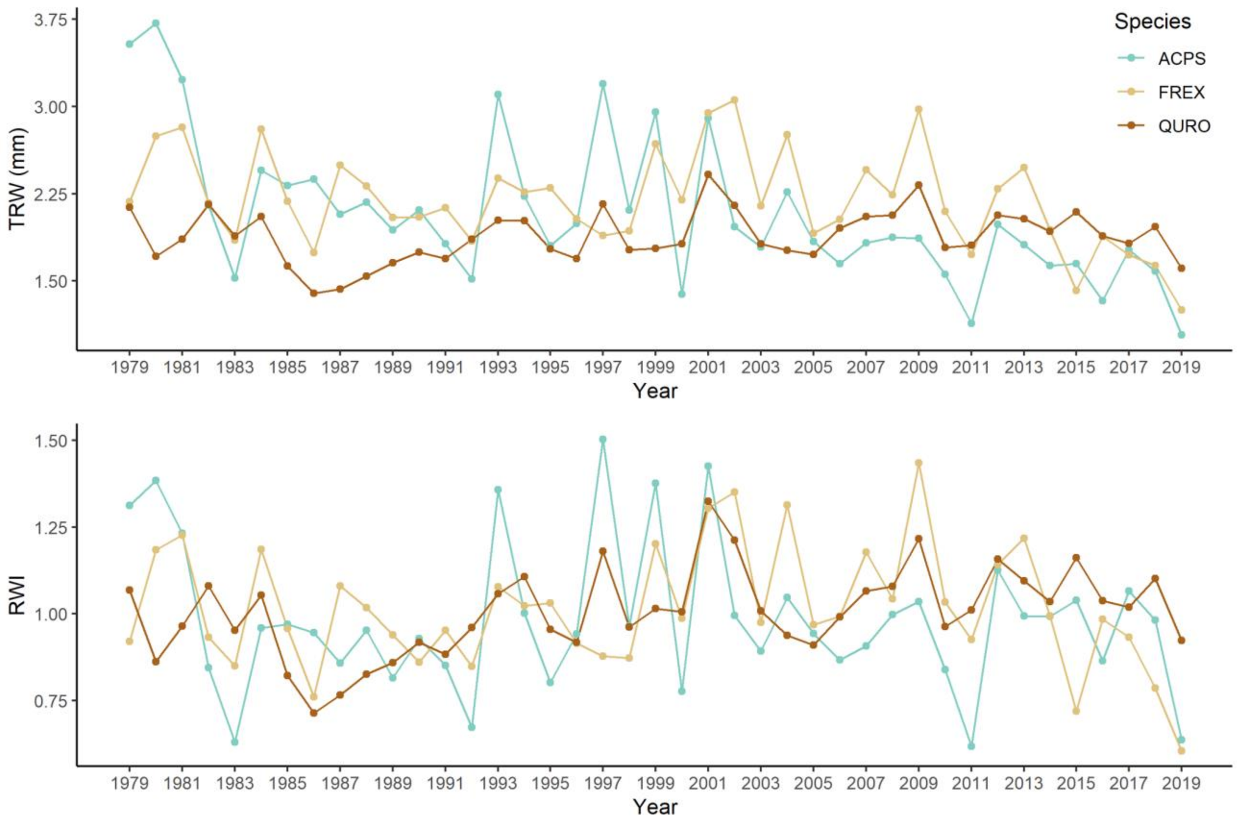


**Fig. S5** Species-specific chronologies of oak (QURO), maple (ACPS) and ash (FREX). Tree ring-width shown as raw ring-width (TRW) and as detrended ring-width index (RWI). Raw and residual chronologies were built using biweight robust means with the dplR package (Bunn, 2008, 2010; Bunn et al., 2020) in R. Ring-widths were detrended with a negative exponential curve. Note that the sites experienced an extreme flood event in January 2011, which likely caused the observed strong growth depression in all species in that year.

**Table S2** Species-specific linear mixed-effects models (LMMs) comparing tree growth responses, stable carbon isotope increases ($\Delta\delta^{13}C$) and drought legacy effects in the consecutive drought years 2018 and 2019 to the average of single drought years.

| Response | Fixed Effects | Estimate | Std.Error | ddf | t | P-value | n |
|---|---|---|---|---|---|---|---|
| **Growth response** | | | | | | | |
| Oak | Intercept | 0.03 | 0.06 | 85 | 0.56 | 0.5763 | 120 |
| | 2018 | 0.09 | 0.06 | 78 | 1.48 | 0.1425 | |
| | **2019** | -0.12 | 0.06 | 78 | -2.00 | **0.0486** | |
| Maple | Intercept | -0.05 | 0.08 | 87 | -0.56 | 0.5744 | 96 |
| | 2018 | 0.14 | 0.11 | 62 | 1.34 | 0.1849 | |
| | **2019** | -0.29 | 0.11 | 62 | -2.74 | **0.0080** | |
| Ash | **Intercept** | -0.18 | 0.03 | 91 | -5.88 | **0.0000** | 126 |
| | **2018** | -0.10 | 0.03 | 82 | -2.94 | **0.0043** | |
| | **2019** | -0.24 | 0.03 | 82 | -7.22 | **0.0000** | |
| **$\Delta\delta^{13}C$** | | | | | | | |
| Oak | **Intercept** | 0.33 | 0.09 | 96 | 3.83 | **0.0002** | 117 |
| | 2018 | 0.02 | 0.10 | 78 | 0.19 | 0.8487 | |
| | **2019** | 0.39 | 0.10 | 78 | 3.93 | **0.0002** | |
| Maple | **Intercept** | 0.41 | 0.15 | 56 | 2.85 | **0.0061** | 78 |
| | 2018 | 0.04 | 0.16 | 50 | 0.26 | 0.7930 | |
| | **2019** | 0.44 | 0.16 | 50 | 2.80 | **0.0074** | |
| Ash | **Intercept** | 0.22 | 0.09 | 84 | 2.50 | **0.0146** | 126 |
| | **2018** | 0.26 | 0.09 | 82 | 2.85 | **0.0056** | |
| | **2019** | 1.34 | 0.09 | 82 | 14.80 | **0.0000** | |
| **Legacy effect** | | | | | | | |
| Oak | Intercept | 0.04 | 0.06 | 67 | 0.72 | 0.4730 | 80 |
| | 2018 | -0.06 | 0.06 | 39 | -0.93 | 0.3580 | |
| Maple | Intercept | 0.01 | 0.07 | 28 | 0.11 | 0.9173 | 64 |
| | **2018** | -0.27 | 0.08 | 49 | -3.52 | **0.0009** | |
| Ash | **Intercept** | 0.14 | 0.04 | 80 | 3.47 | **0.0008** | 84 |
| | **2018** | -0.43 | 0.05 | 41 | -8.41 | **0.0000** | |

Note: Significant fixed effects ($P<0.05$) printed in bold. Shown are species-specific, linear mixed-effects models (LMMs) fit with the packages lme4 (Bates, Mächler, Bolker, & Walker, 2015) and lmerTest (Kuznetsova, Brockhoff, & Christensen, 2017) in R, using a significance level of $\alpha=0.05$ and restricted maximum likelihood estimation (REML). For growth responses and $\Delta\delta^{13}C$ we present here the most parsimonious models selected based on backward model selection (see Methods for details) that contain drought event as fixed effect and tree identities nested within plot as random effect. The initial full model contained drought event (single droughts, 2018, 2019), groundwater level (dry > 2m, moist ≤ 1m) and their interaction as fixed effects (see Methods for details) but groundwater level as well as the interaction between drought event and groundwater level were, if not significant ($P>0.05$), dropped during model selection based on F-tests using Satterthwaite degrees of freedom (see Kuznetsova et al. (2017) for details). Drought events were fitted as categorical fixed effects, i.e. the fixed effects test for significant difference in the growth response and $\Delta\delta^{13}C$ in 2018 and 2019 compared to the intercept, that is, former single droughts (using the mean growth response and $\Delta\delta^{13}C$ in the drought years 2003, 2006 and 2015). We analysed drought legacy effects using the same modelling procedure. Specifically, we used LMMs with drought event (mean of single droughts and 2018) as fixed effects and tree identity nested within plot as random effect (except for the drought legacy LMM of maple which was most parsimonious with plot as the only random effect). All analyses were conducted in R version 4.1.0 (R Core Team, 2021).

**Table S3** Pairwise comparisons of the growth responses and stable carbon isotope increases ($\Delta\delta^{13}C$) between the average of single drought years and the consecutive drought years 2018 and 2019.

| Species | Single drought vs 2018 | Single drought vs 2019 | 2018 vs 2019 |
|---|---|---|---|
| **Growth response** | | | |
| Oak | 0.3054 | 0.1182 | **0.0023** |
| Maple | 0.3783 | **0.0214** | **0.0004** |
| Ash | **0.0118** | **0.0000** | **0.0001** |
| **$\Delta\delta^{13}C$** | | | |
| Oak | 0.9805 | **0.0006** | **0.0012** |
| Maple | 0.9622 | **0.0198** | **0.0381** |
| Ash | **0.0153** | **0.0000** | **0.0000** |

Note: Significant fixed effects ($P<0.05$) printed in bold. Post-hoc tests were performed for the models presented in Supplementary Table 1 with the emmeans package (Lenth, 2020) with p-values adjusted for multiple comparisons using Tukey's Honest Significant Difference.

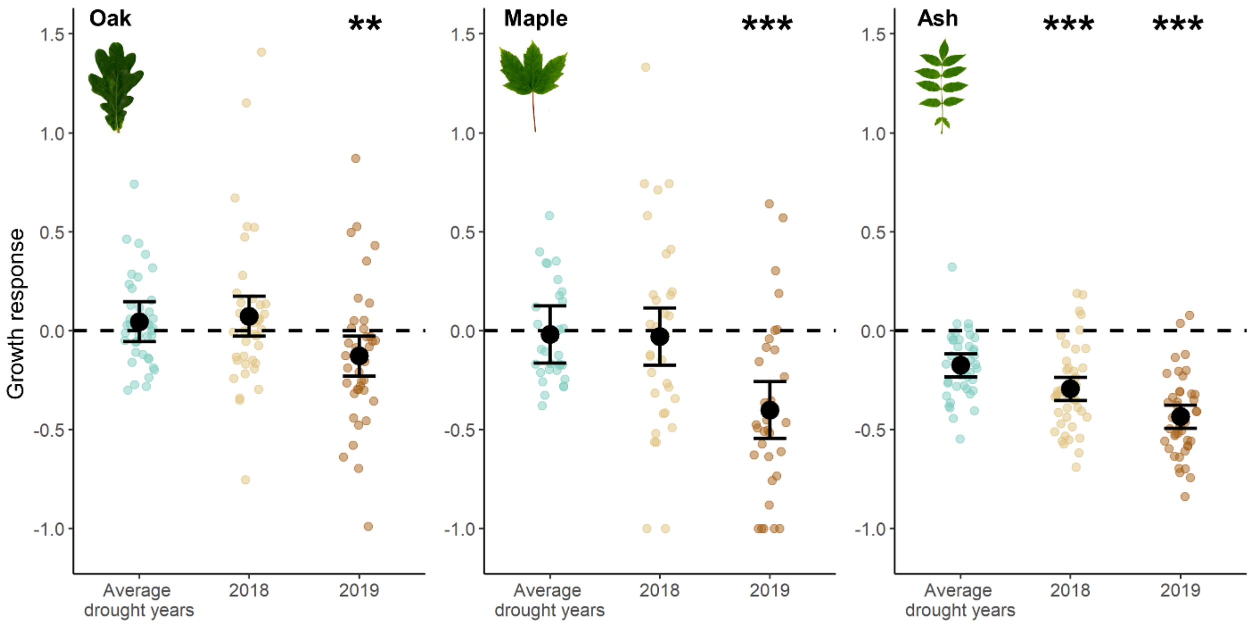


**Fig. S6** Growth response of oak, maple and ash trees in drought years calculated based on raw instead of detrended tree ring-widths. The figure shows the growth response in the consecutive hotter drought years 2018 and 2019 compared to the mean growth response in single drought years (2003, 2006 and 2015). Zero corresponds to a comparable growth in dry and normal years. Black points show estimated marginal means and error bars the 95% confidence intervals of linear mixed-effects model fits. Coloured points show growth response values per tree and species (oak n=40, maple n=32, ash n=42) and are jittered to enhance visibility. The growth response was calculated as the annual ring-width in drought year(s) / average annual ring-width in normal climatic years (2005, 2009, 2017; Eq. 1). Statistically significant differences in the growth response between the years 2018 and 2019 compared to single drought years are indicated by asterisks over the respective year ('***' p <0.001; '**' p <0.01; '* 'p <0.05).

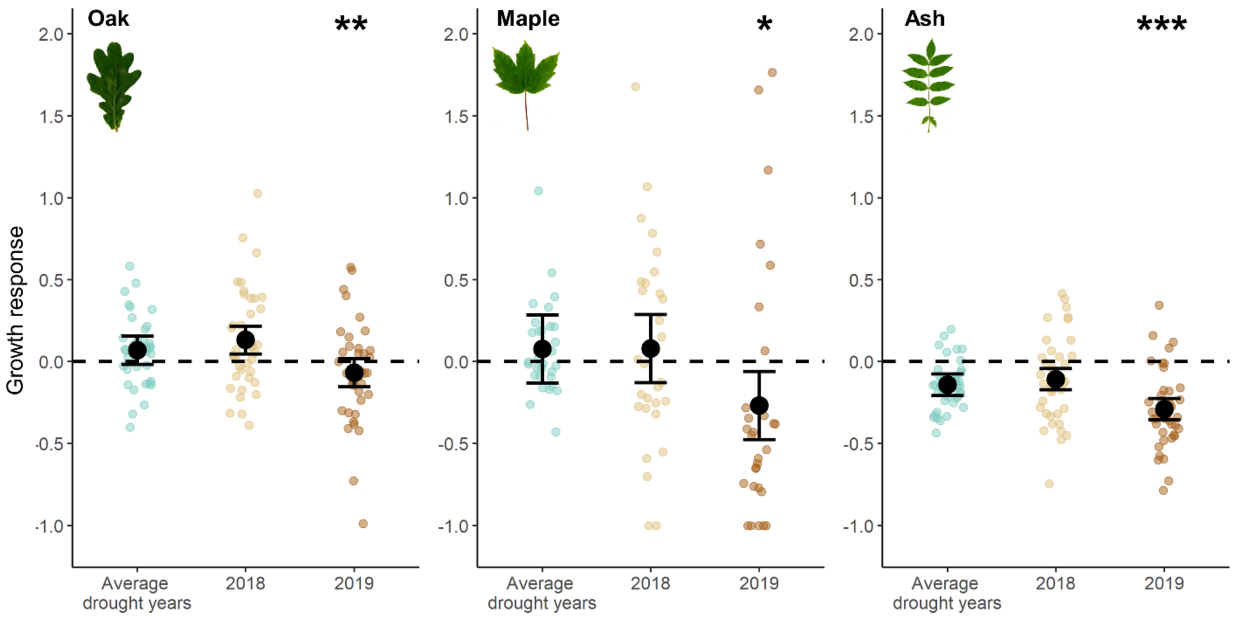


**Fig. S7** Growth response of oak, maple and ash in drought years calculated based on a one-year reference period. The figure shows the growth response in the consecutive hotter drought years 2018 and 2019 compared to the mean growth response in single drought years (2003, 2006 and 2015). Zero corresponds to a comparable growth in dry and normal years. Black points show estimated marginal means and error bars the 95% confidence intervals of linear mixed-effects model fits. Coloured points show growth response values per tree and species (oak n=40, maple n=30, ash n=42) and are jittered to enhance visibility. The growth response was calculated as the annual ring-width in drought year(s) / ring-width in the only climatically normal year (2017) preceding the 2018–2019 consecutive drought with Eq. 1. Note that 2015 but also 2016 were characterized by exceptionally dry conditions (Supplementary Fig. 1), which precludes their use as reference years. We also excluded two maple trees with exceptionally high growth response values (3.4 and 5.3) from this analysis, which were caused by their very low growth in 2017, probably due to waterlogging. This exclusion did not effect the results but highlights the merit of using several reference years instead of a single year to avoid such outliers. The tree ring-widths have been detrended with a negative exponential function. Statistically significant differences in the growth response between the years 2018 and 2019 compared to single drought years are indicated by asterisks over the respective year ('***' p <0.001; '**' p <0.01; '* 'p <0.05).

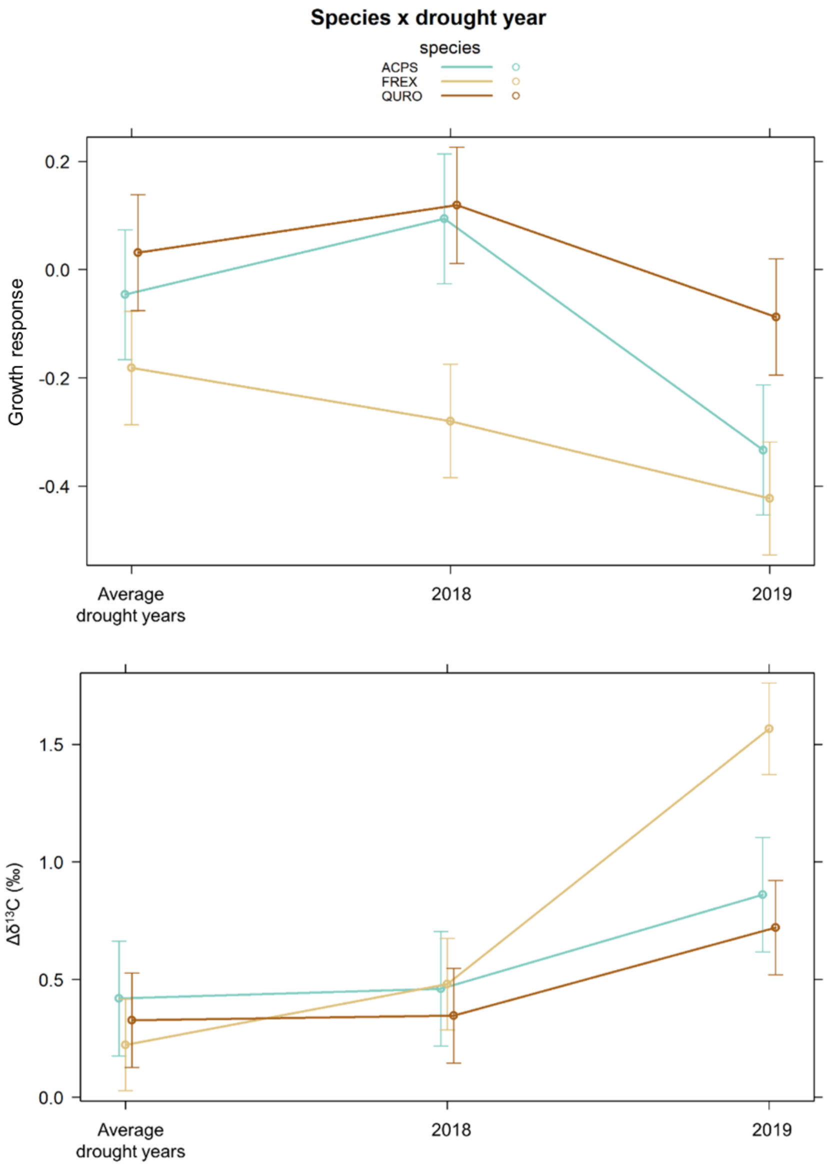


**Fig. S8** Growth response and increase in the carbon isotope ratio ($\Delta\delta^{13}C$) in wood of oak, maple and ash in drought years. The figure shows the growth response (upper panel) and $\Delta\delta^{13}C$ (lower panel) in the consecutive hotter drought years 2018 and 2019 compared to the mean growth response and $\Delta\delta^{13}C$ in single drought years (2003, 2006 and 2015). In contrast to the species-specific models presented in manuscript Fig. 1 here presented estimated marginal means and error bars (95% confidence intervals) are based on linear mixed-effects models that predict the growth response and $\Delta\delta^{13}C$ by a significant interaction between drought year and species identity, with P=0.0105 for the growth response and P<0.0001 for $\Delta\delta^{13}C$, respectively. All other model specifications (including random effect structure and R packages used) are the same. Sample size is n=114 trees for the growth response and n=107 for $\Delta\delta^{13}C$, that is, the sum of the sample sizes of oak, maple and ash. Both models show the same response pattern that we obtain from the species-specific models (Fig. 2).

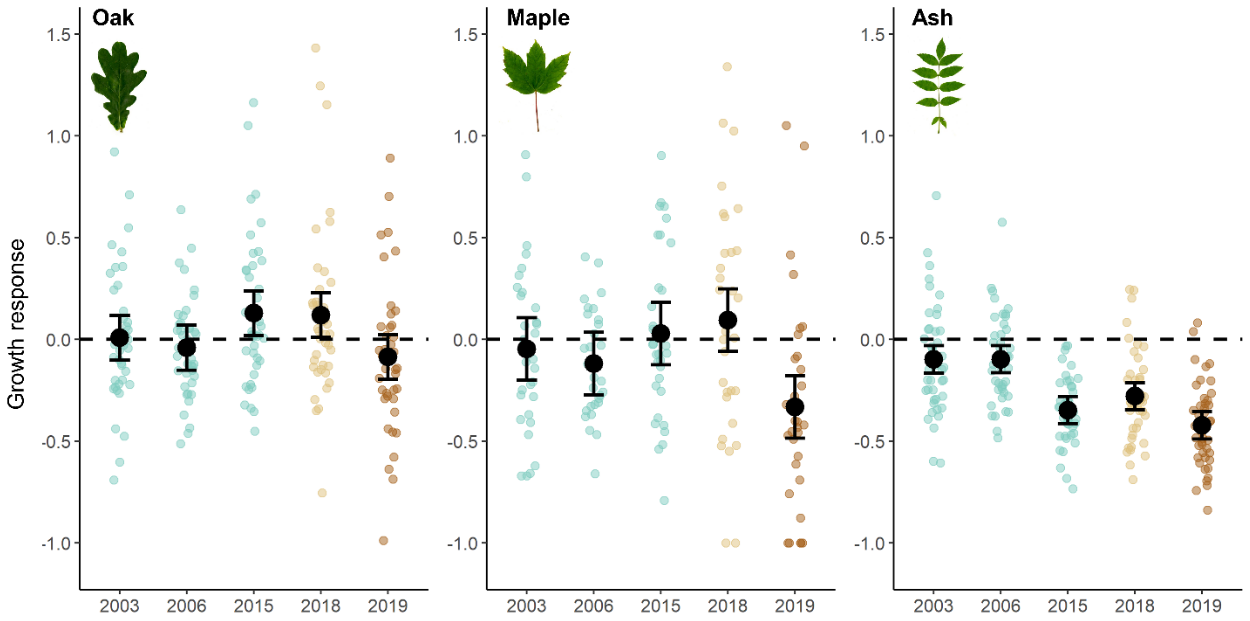


**Fig. S9** Growth response of oak, maple and ash in drought years. The figure shows the growth response in the consecutive hotter drought years 2018 and 2019 compared to the growth response in each single drought year (2003, 2006 and 2015). Zero corresponds to a comparable growth in dry and normal years. Black points show estimated marginal means and error bars the 95% confidence intervals of linear mixed-effects model fits. Coloured points show growth response values per tree and species (oak n=40, maple n=30, ash n=42) and are jittered to enhance visibility. The growth response was calculated with Eq. 1. The tree ring-widths have been detrended with a negative exponential function. The shown models have no baseline, such as the mean growth response across single drought years (Fig. 3), against which the effects of the consecutive drought years 2018 and 2019 can be compared. We therefore tested for statistically significant differences between growth responses in each examined drought year using post-hoc pairwise comparisons (Table S4). The shown models yielded overall similar conclusions compared to the baseline models shown in the main manuscript (Fig. 3). The hotter drought in 2018 did not induce growth responses in oak and maple that differed significantly from growth responses in any single drought year (Table S4). In contrast, growth reductions in 2019, the second consecutive drought year, were significantly stronger than in the preceding drought year 2018 for all species (Table S4). Only for ash, growth responses were significantly stronger in 2018 compared to 2003 and 2006 (Table S4), but the growth response in 2015 was intermediate between responses observed in 2018 and 2019 (Fig. S9). This stronger response in 2015 may be related to incipient ash-dieback causing reduced growth in this species in addition to the effects of drought as ash-dieback arrived in the examined stands only in 2011 (first reports of ash-dieback by the local forest administration). Nonetheless, ash responded with reduced growth in all examined single drought years (Fig. S9) in line with our conclusion of generally strongest growth responses in this species (Fig. 3).

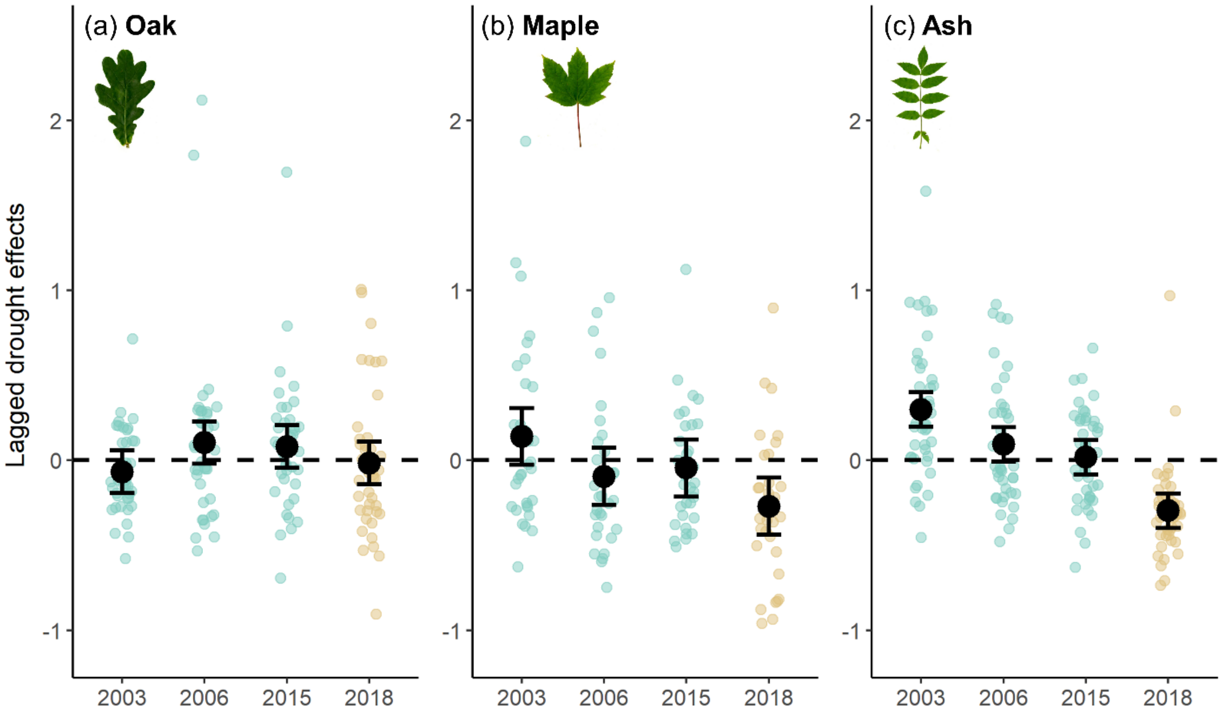


**Fig. S10** Drought legacy effects in growth of oak, maple and ash in the year following drought events. The figure shows drought legacy effects induced by the hotter drought year 2018 compared to legacy effects induced by each single drought year (2003, 2006 and 2015). Legacy effects were quantified as observed minus predicted (detrended) tree ring-width based on climate in year 1 after the drought event. Zero corresponds to growth as expected based on climate conditions, while negative values indicate drought legacies in form of lower than expected post-drought growth. Black points show estimated marginal means and error bars the 95% confidence intervals of linear mixed-effects model fits. Coloured points show legacy effects per tree and species (oak n=40; maple n=32; ash n=42) and are jittered to enhance visibility. The shown models have no baseline, such as the mean legacy effect across single drought years (Fig. 4), against which the effects of the consecutive drought years 2018 and 2019 can be compared. We therefore tested for statistically significant differences between legacy effects in each examined drought year using post-hoc pairwise comparisons (Table S4). The shown models yielded overall similar conclusions compared to the baseline models shown in the main manuscript (Fig. 4). Observed tree growth in year 1 after any of the single droughts was not lower than growth predicted based on climate (Fig. S10). In contrast, the hotter drought year 2018 induced substantial legacy effects in maple and ash but not in oak.

**Table S4** Pairwise comparisons of the growth responses and drought legacy effects between each single drought year and the consecutive drought years 2018 and 2019.

| Species | 2003 vs 2006 | 2003 vs 2015 | 2003 vs 2018 | 2003 vs 2019 | 2006 vs 2015 | 2006 vs 2018 | 2006 vs 2019 | 2015 vs 2018 | 2015 vs 2019 | 2018 vs 2019 |
|---|---|---|---|---|---|---|---|---|---|---|
| **Growth response** | | | | | | | | | | |
| Oak | 0.9573 | 0.4045 | 0.4825 | 0.6477 | 0.1081 | 0.1435 | 0.9615 | 0.9999 | **0.0179** | **0.0260** |
| Maple | 0.9580 | 0.9446 | 0.6393 | **0.0494** | 0.6033 | 0.2374 | 0.2347 | 0.9690 | **0.0054** | **0.0006** |
| Ash | 1.0000 | **0.0001** | **0.0002** | **0.0000** | **0.0000** | **0.0002** | **0.0000** | 0.4728 | 0.3898 | **0.0070** |
| **Legacy effect** | | | | | | | | | | |
| Oak | 0.1707 | 0.2830 | 0.9233 | - | 0.9927 | 0.4781 | - | 0.6507 | - | - |
| Maple | 0.1026 | 0.2687 | **0.0006** | - | 0.9622 | 0.3190 | - | 0.1285 | - | - |
| Ash | **0.0182** | **0.0005** | **0.0001** | - | 0.6928 | **0.0001** | - | **0.0001** | - | - |

Note: Significant fixed effects ($P<0.05$) printed in bold. Post-hoc tests were performed for the models presented in Supplementary Fig. S9–S10 with the emmeans package (Lenth, 2020) with p-values adjusted for multiple comparisons using Tukey's Honest Significant Difference.

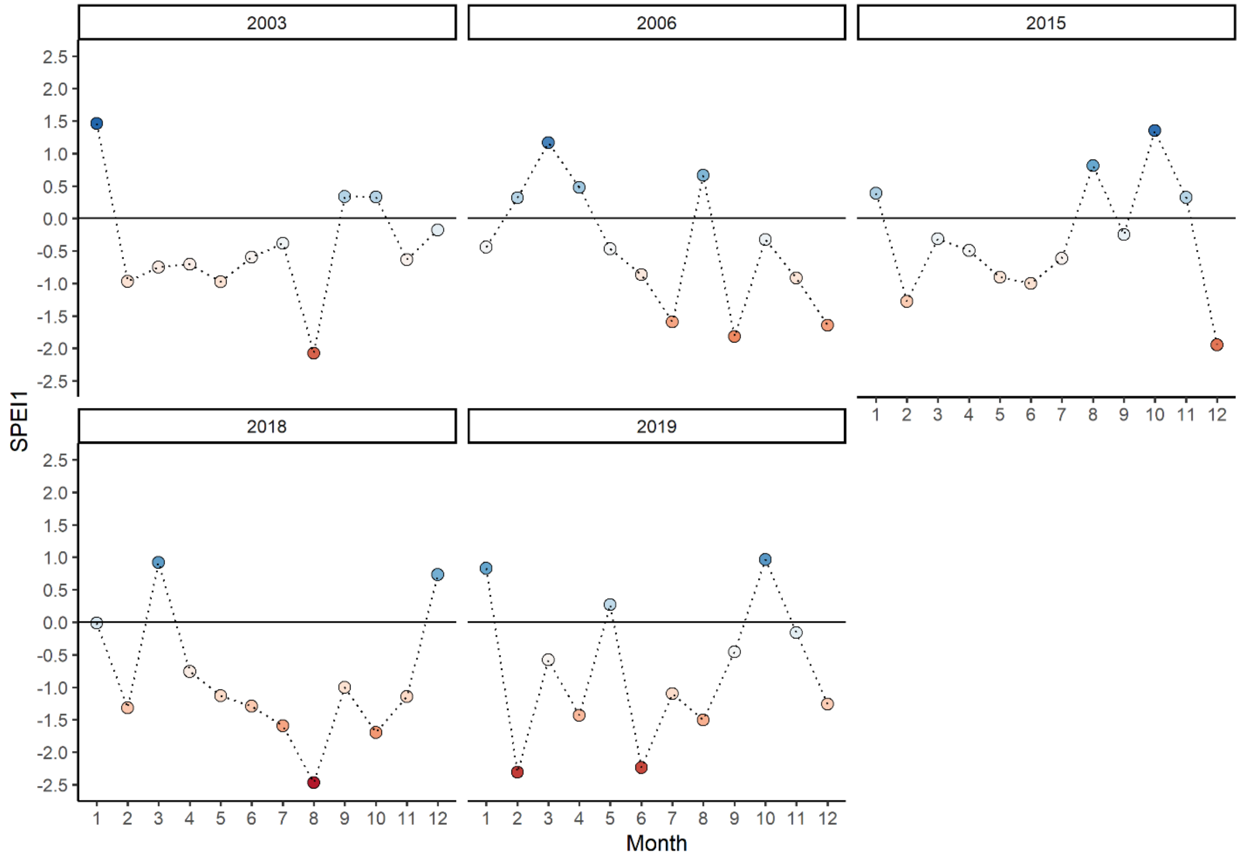


**Fig. S11** Drought onset. Shown are monthly standardized water balances of precipitation minus potential evapotranspiration (A–C, Standardized Precipitation Evapotranspiration Index (SPEI)) for the examined single drought years (2003, 2006, 2015, upper panels) as well as for the consecutive drought years 2018 and 2019 (lower panels) in the Leipzig floodplain forest. Points are colored according to their value with deeper red indicating increasing drought severity. SPEI values for each drought year are shown for each month as SPEI1, i.e. as standardized water balance in that month. The horizontal line in (A–C) represents the long-term mean, negative values indicate water deficits and positive values water surpluses. SPEI values below -1 and above 1 can be considered as exceptionally dry and wet, respectively (McKee et al., 1993), that is, as indicative of drought onset. See Supplementary Fig. 1 for further details on the SPEI calculation.

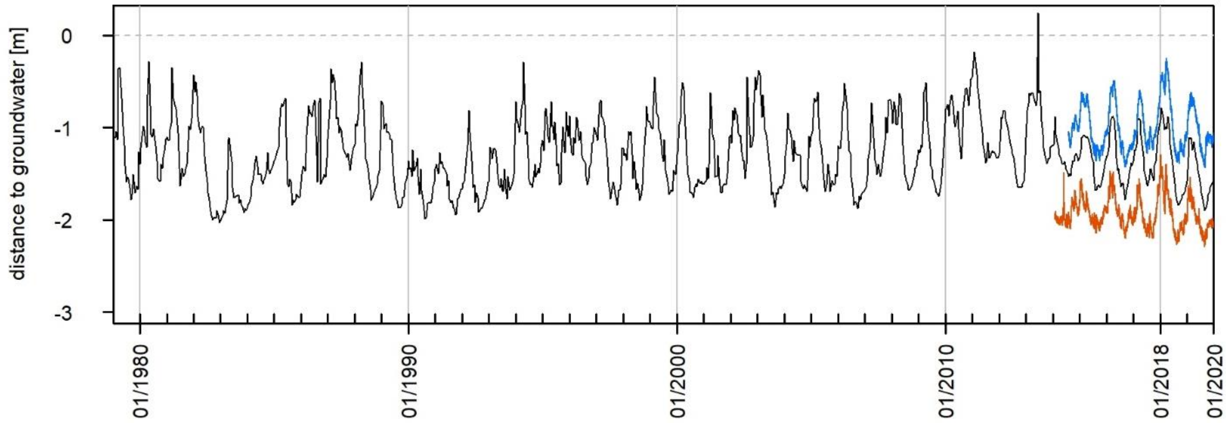


**Fig. S12** Groundwater levels (shown as distance to groundwater below soil surface) in the Leipzig floodplain forest. The black line shows groundwater levels over the last 40 years (1979–2019) at the official gauging station (46390103 Schkeuditz, LfULG 2021). The colored lines show groundwater levels for two representative plots of the sampled plot network (see Supplementary Fig. 6) with the blue line showing groundwater levels at a moist (mean distance to groundwater ≤ 1m) and the red line groundwater levels at a dry plot (mean distance to groundwater > 2m), respectively. The dashed line represents the soil surface. Groundwater measurements of the Lebendige Luppe project were only available for the last six years. During the last 40 years we did not observe a long-term trend (neither decrease nor increase) of groundwater levels. During the consecutive drought years 2018–2019, we observed no trend in groundwater levels on moist plots but the lowest summer minima of the observation period on dry plots. However, absolute changes in groundwater level were small and are thus unlikely a major driver of observed tree drought responses (as also indicated by the overall small effect of groundwater level on tree growth and $\Delta\delta^{13}C$). The small change in groundwater level during the 2018–2019 drought are consistent with relatively low changes in river discharge (Supplementary Fig. 2).

### References


Bates, D., Mächler, M., Bolker, B., & Walker, S. (2015). Fitting Linear Mixed-Effects Models Using lme4. *Journal of Statistical Software*, *67*(1). https://doi.org/10.18637/jss.v067.i01

Beguería, S., & Vicente-Serrano, S. M. (2017). SPEI: Calculation of the Standardised Precipitation-Evapotranspiration Index. [Computer software]. Retrieved from https://CRAN.R-project.org/package=SPEI

Bunn, A. G. (2008). A dendrochronology program library in R (dplR). *Dendrochronologia*, *26*(2), 115–124. https://doi.org/10.1016/j.dendro.2008.01.002

Bunn, A. G. (2010). Statistical and visual crossdating in R using the dplR library. *Dendrochronologia*, *28*(4), 251–258. https://doi.org/10.1016/j.dendro.2009.12.001

Bunn, A. G., Korpela, M., Biondi, F., Campelo, F., Mérian, P., Qeadan, F., & Zang, C. (2020). *dplR: Dendrochronology Program Library in R*. Retrieved from https://CRAN.R-project.org/package=dplR

DWD Climate Data Center (CDC). Monthly mean of station observations of precipitation in mm, air temperature at 2 m above ground in °C, daily air temperature minimum at 2 m above ground in °C, daily air temperature maximum at 2 m above ground in °C, wind force in bft, cloud coverage in eighth, air pressure at station level in hPa, relative humidity at 2 m above ground in %, vapor pressure in hPa for Germany, version v19.3, last accessed: 06.02.2020. Data for Station Leipzig/Halle, ID 2932.

Granier, A. (1987). Evaluation of transpiration in a Douglas-fir stand by means of sap flow measurements. *Tree Physiology*, *3*(4), 309–320. https://doi.org/10.1093/treephys/3.4.309

Kuznetsova, A., Brockhoff, P. B., & Christensen, R. H. B. (2017). lmerTest Package: Tests in Linear Mixed Effects Models. *Journal of Statistical Software*, *82*(13). https://doi.org/10.18637/jss.v082.i13

Lenth, R. (2020). emmeans: Estimated Marginal Means, aka Least-Squares Means. [Computer software]. Retrieved from https://CRAN.R-project.org/package=emmeans

McKee, T. B., Doesken, N. J., & Kleist, J. (1993). The relationship of drought frequency and duration to time scales. In *Proceedings of the 8th Conference on Applied Climatology*.

R Core Team (2021). A language and environment for statistical computing [Computer software]. Vienna, Austria: R Foundation for Statistical Computing: R Foundation for Statistical Computing. Retrieved from https://www.R-project.org/

Scholz, M., Seele, C., Engelmann, R. A., Hartmann, H., Heinrich, J. Henle, K., Herkelrath, A., . . . Riedel, J. (2018). Das Projekt Lebendige Luppe – ein Beitrag zur Renaturierung der Leipziger Nord-West-Aue. *Auenmagazin*, *14*(14), 14–21.

Weil, R. R., & Brady, N. C. (2017). *Nature and properties of soils* (15th edition). Pearson.

Zang, C., & Biondi, F. (2015). treeclim: an R package for the numerical calibration of proxy-climate relationships. *Ecography*, *38*(4), 431–436. https://doi.org/10.1111/ecog.01335
